# iSEE: Interface Structure, Evolution and Energy-based machine learning predictor of binding affinity changes upon mutations

**DOI:** 10.1101/331280

**Authors:** Cunliang Geng, Anna Vangone, Gert E. Folkers, Li C. Xue, Alexandre M.J.J. Bonvina

## Abstract

Quantitative evaluation of binding affinity changes upon mutations is crucial for protein engineering and drug design. Machine learning-based methods are gaining increasing momentum in this field. Due to the limited number of experimental data, using a small number of sensitive predictive features is vital to the generalization and robustness of such machine learning methods. Here we introduce a fast and reliable predictor of binding affinity changes upon single point mutation, based on a random forest approach. Our method, iSEE, uses a limited number of **i**nterface **S**tructure, **E**volution and **E**nergy-based features for the prediction. iSEE achieves, using only 31 features, a high prediction performance with a Pearson correlation coefficient (PCC) of 0.80 and a root mean square error of 1.41 kcal mol^-1^ on a diverse training dataset consisting of 1102 mutations in 57 protein-protein complexes. It competes with existing state-of-the-art methods on two blind test datasets. Predictions for a new dataset of 540 mutations in 58 protein complexes from the recently published SKEMPI 2.0 database reveals that none of the current methods perform well (PCC<0.4), although their combination does improve the predictions. Feature analysis for iSEE underlines the significance of evolutionary conservations for quantitative prediction of mutation effects.

## Introduction

The affinity between proteins and their binding partners is a fundamental property that governs their function in cells. Mutations in proteins can induce changes in the binding affinity for their interaction partners, altering their functioning by perturbing their communication network. Missense mutations are often linked to various human diseases^1^, such as cancer. Quantitative characterization of binding affinity changes can therefore shed light on the relation between coding variations and disease phenotypes, and guide the design of effective therapeutics for genetic disorders. It can also be particularly useful for engineering protein-protein interactions with modulated binding affinity.

Various experimental methods can be used to quantitatively measure binding affinities^2,3^, each with their own limitations and precision. Although they provide valuable information, experimental methods can be labor-intensive and time-consuming, and, as a consequence, lag behind the rapid advances of sequencing technologies, which are generating a huge amount of data on disease-causing mutations. This calls for the development of reliable and fast computational methods for estimating the mutation effects on binding affinity (i.e., the binding free energy change between a wild type and mutant complex, ΔΔG).

Computational methods for ΔΔG prediction can be largely grouped into three main strategies:

1) Rigorous methods, such as thermodynamic integration and free energy perturbation^4,5^, 2) empirical energy-based methods, based for example on classical mechanics or statistical potentials^6-10^ (typically in linear forms), and 3) machine learning-based methods which can exploit a large variety of energetics and non-energetics (e.g. geometric, evolutionary) features^11-13^. The rigorous methods can be accurate but they are computationally highly demanding. Their application is, therefore, limited to mainly low-throughput and small system ΔΔG calculations. The empirical energy-based methods are much faster and more broadly applied. They usually take a form of linear functions, often with only energy-based terms, and fail to exploit evolution information, which can limit their ability to capture mutation effects on binding affinity. Insufficient conformational sampling, especially for mutations in flexible regions, can limit the accuracy of energy-based methods. In contrast, machine learning-based methods are potentially less sensitive to this since they can model mutation effects using not only potentials or energies but also other relevant features, such as, sequence, structure and evolution. Machine learning approaches typically aim to model the intrinsic relationship between features of a mutation site and the response variable (e.g., the binding affinity change) by training statistical models from mutation datasets with experimentally determined ΔΔG. Due to the data-driven essence of machine learning, the availability of a large amount of reliable experimental data and the construction of features that can reflect structural and physico-chemical changes caused by mutations are crucial factors for the success for this type of methods. It is therefore not surprising that the publication of the SKEMPI database^14^ (version 1.1, which was in the past six years the largest mutation ΔΔG dataset for protein-protein complexes containing 3047 mutations in 85 complexes) quickly promoted the emergence of several machine learning-based ΔΔG predictors^11-13^. However, the SKEMPI 1.1 dataset is still rather limited in size and one has to be careful not to use too many features to train a model to avoid overfitting problems. It is therefore important to design fast and reliable ΔΔG predictors that exploit only a limited number of sensitive and relevant features. Very recently, an update of SKEMPI was published, version 2.0^15^, which provides a much extensive dataset and gives us the opportunity to test various predictors on data none of them has previously seen.

Residue conservation plays a central role in determining the binding affinity. It has been verified that the binding energy is not evenly distributed among the interfacial residues. Instead, some residues (hot-spots) contribute most to the binding affinity^16-18^. Such residues are often highly conserved. Interface conversation has been used in several of the best performing ΔΔG predictors^13,19^. However, since the conservation measure they used is structure-based, relying on the availability of structural homologs^13,19^, the application and prediction of these ΔΔG predictors are largely limited by the availability and the number of such homologs. By contrast, conservation from Position Specific Scoring Matrix (PSSM) is sequence-based and thus better applicable. The PSSM value is a log likelihood ratio between the observed probability of one type of amino acid appearing in a specific position in the multiple sequence alignment (MSA) and the expected probability of that amino acid type appearing in a random sequence^20^. Thus, each position of a protein can be represented as a 20 by 1 PSSM profile (or vector), which captures the conservation property of each amino acid type at a specific position.

Here we present a machine learning-based method named iSEE (interface Structure, Evolution and Energy-based ΔΔG predictor), which combines HADDOCK^21^, structure and energy terms of wildtype and mutant complexes as well as PSSM conservation profiles before and after mutations (**Figure 1**). HADDOCK^21^ is our in-house docking program, which has been consistently ranking among the top predictors and scorers in CAPRI, a community-wide experiment for the prediction of biomolecular interactions^22^. Its simple but sensible scoring function has contributed much to its success^23^. It includes intermolecular van der Waals (Evdw) and Coulomb electrostatics (Eelec) energies, an empirical desolvation energy term (Edesolv)^24^ and buried surface area (BSA), which is only used in intermediate scoring steps and not in the final scoring function. iSEE is based on a random forest model^25^ for ΔΔG prediction, trained on a subset of 1102 single point mutations from 57 complexes from SKEMPI 1.1. It uses a small number of features (31) to lower the overfitting risk and competes with both empirical potentials- and machine learning-based state-of-the-art methods on an existing independent test dataset (the Benedix *et al.* NM dataset^8^) and a large test dataset from the recently released SKEMPI 2.0 database. The recent release of SKEMPI 2.0 allows us for the first time to test various ΔΔG predictors on a large new blind dataset with more than 500 mutations. Analysis of the importance of the features used in iSEE highlights the significance of evolutionary information in predicting the effect of mutations on the binding affinity of protein complexes.

**Figure 1.**
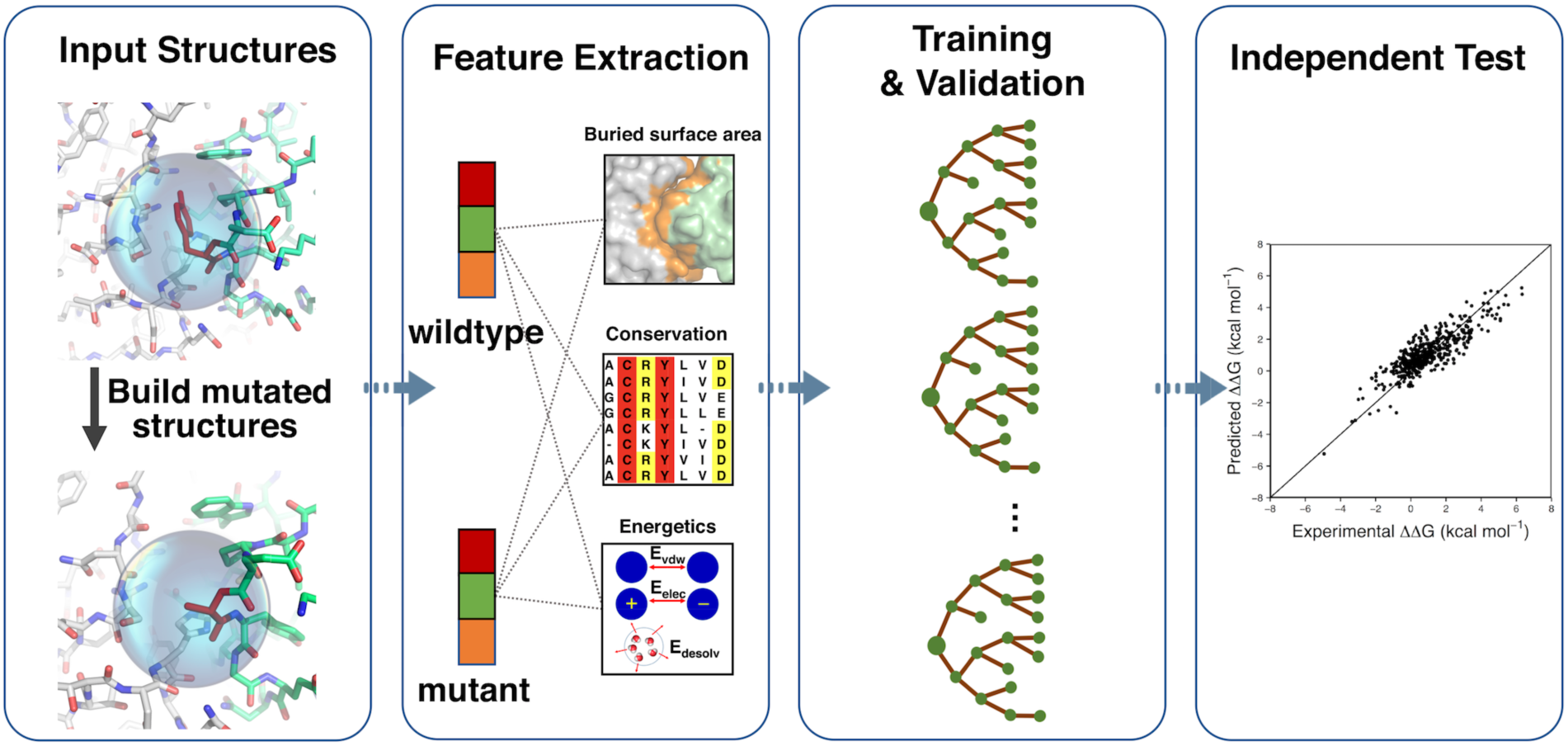
The workflow of iSEE predictor. Only the 3D structure of wildtype complex and mutation information are necessary input for iSEE. We first model the mutated structure using HADDOCK (the water refinement web service). Then we extract features related to the evolutionary conservation and to changes in structure and energetics caused by the mutation. A random forest algorithm is then optimized and cross validated on a training dataset, resulting in our final ΔΔG predictor iSEE. Finally, iSEE is evaluated on two blind test datasets and compared with other current leading ΔΔG predictors.

## Methods

### Training and test datasets

Three datasets of experimental ΔΔG with available crystal structures of protein complexes were used in this study. Only single point mutations in the interface of the protein-protein complexes were considered, and only for dimeric complexes for ease of computations, but our prediction scheme can be easily extended to multimers. The interface residues were defined following Levy’s method^26^ as those located in the core, rim and support regions.

The training dataset was extracted from the DACUM database (https://github.com/haddocking/DACUM)^27^, our ΔΔG database derived from the SKEMPI 1.1 database^14^. DACUM contains 1872 single point mutations in 81 protein complexes. After applying the above-mentioned filter criteria, 1102 single point mutations in 57 protein complexes were selected.

We compiled two independent datasets, not used for training, to evaluate and compare our predictor with state-of-the-art ΔΔG predictors.

We selected a subset from the Benedix *et al.* NM dataset^8^ for which predictions of various ΔΔG predictors have already been reported^7,9^. The original NM dataset has both single point and multiple point mutations in protein dimer or multimer complexes. We applied the same filtering criteria as above. Moreover, to avoid any overlap between the training dataset and the test dataset, mutations existing in the training dataset were filtered out from the original NM dataset. This procedure resulted in 19 mutations in one complex (PDB ID: **1IAR**). For this heterodimer complex, only ΔΔG values for mutations on chain A were contained in our training dataset, while all ΔΔG data in the NM dataset are for mutations on chain B. Thus, the NM dataset is distinct from our training dataset at the level of mutation position. In the remaining of our paper, we will refer to those data as “the NM dataset”.

Finally, we selected new data from the recently released SKEMPI 2.0 database. After applying the above-mentioned filter criteria, 540 mutations in 58 protein complexes were selected. We will refer to this dataset as “the S540 dataset” in the remaining paper.

### Predictive features

We compiled a list of 31 features including intermolecular energy terms and buried surface area (BSA) from HADDOCK^21^ and conservation values from PSSM.

To obtain the structural and energetic features, both wild type and mutant structures were refined using the protocol implemented in the refinement interface of the HADDOCK server^28^. The mutations were introduced by simply changing the identity of the residue in the coordinate file and letting HADDOCK rebuild the missing side-chain atoms and refine the interface in explicit solvent using the TIP3P water model and the Optimized Potentials for Liquid Simulations (OPLS) force field^29^ with an 8.5Å cutoff for the non-bonded interactions. The HADDOCK terms for wildtype and mutant complex were extracted from the top ranked HADDOCK model. The HADDOCK-derived features are:

- Evdw, the intermolecular van der Waals energy described by a 12-6 Lennard-Jones potential.
- Eelec, the intermolecular electrostatic energy described by a Coulomb potential.
- Edesolv, an empirical desolvation energy term^24^.
- BSA, the buried surface area calculated by taking the difference between the sum of the solvent accessible surface area (SASA) for each individual protein and the SASA of the protein complex using 1.4Å water probe radius.

The four HADDOCK terms of wildtype complex and the differences of the HADDOCK terms between mutant and wildtype complexes were used as HADDOCK-based predictive features, which are named as Evdw_wt, Eelec_wt, Edesolv_wt, BSA_wt, Evdw_diff, Eelec_diff, Edesolv_diff and BSA_diff.

The PSSM was calculated through PSI-BLAST of BLAST 2.3.0^30^ using a local version of the software and databases with the following parameters: BLOSUM62 was used as scoring matrix by default, and PAM30 was used when BLOSUM62 failed for short sequences; the number of iterations was 3 and the e-value threshold was set to 0.0001; the BLAST database was the nr database (non-redundant BLAST curated protein sequence database, version on Aug. 22^nd^ 2016). Default values were used for all the other parameters. For each mutation position of a query protein, 4 types of conservation features were extracted from the PSSM file:

- the PSSM profile for this position, which is a 20 by 1 vector (PSSM_AA).
- the information content for this position (PSSM_IC).
- the individual PSSM value for the wildtype residue at this position (PSSM_wt).
- the difference between the individual PSSM values for mutant residue and wildtype residue at this position (PSSM_diff).

### Training procedures and evaluation metrics

We used the random forest algorithm^31^ from the R Caret package^32^ to train our ΔΔG predictor. We optimized the parameters of random forest over 10 times 10-fold cross-validations on the training dataset: the number of trees to grow, defined by the “ntree” parameter, was varied from 10 to 100 in steps of 10, and the number of variables randomly sampled as candidates at each split, defined the “mtry” parameter, was sampled from 1 to 20. The prediction performance was evaluated by root mean square error (RMSE) and Pearson’s correlation coefficient (PCC).

### Comparison with other ΔΔG predictors

The performance of the iSEE ΔΔG predictor was compared with several state-of-the-art ΔΔG predictors on the independent NM and SKEMPI 2.0 S540 test datasets. For the NM dataset, the predicted ΔΔG values of pred1^9^, pred2^9^, CC/PBSA^8^, BeAtMuSiC^10^ and FoldX^6^ were directly extracted from Li et al^9^ and those of ZeMu from Dourado’s paper^7^. Predictions of mCSM^11^ and BindProfX^19^ for the test datasets were directly obtained from their respective webservers. The default parameters of BindProfX were used except the “Score to use” which was set to “interface profile and physics potential” (the authors reported it to work best for single point mutations^19^). A local version of FoldX (4.0) was used for the S540 dataset.

### Classification of mutations

Mutation were classified based on three scenarios: the location of the mutation, the type of mutated amino acid and the change in the size of the amino acid side-chain.

Based on the type of secondary structure a mutation is located, it was classified as a loop or non-loop mutation. We used DSSP^33,34^ (v2.0.4) for secondary structure assignment. DSSP code S, B and blank were considered as loop, otherwise non-loop.

Based on the type of mutated amino acid, a mutation was called “toALA” mutation when a residue was mutated to alanine, otherwise “toNonALA” mutation.

The change of amino acid size was defined as the difference of volumes (ΔV) between mutant and wildtype amino acids. The volumes of the 20 amino acids were taken from literature^35^. A mutation was classified as “neutral” if | ΔV | ≤ 10 Å^3^, as small to large (“toLarge”) if ΔV > 10 Å^3^, and as large to small (“toSmall”) if ΔV < −10 Å^3^.

### Feature importance analysis

We used the algorithm from the R package randomForest^31^ to evaluate feature importance. The feature importance is measured by the decrease of mean squared error when splitting on a feature, averaged over all trees. The importance measure of a group of features was calculated by taking the sum of weighted importance of each feature in that group with the weight for each feature defined as the number of times the feature was chosen as split variable over all trees divided by the total of all group member features. The PSSM profile scores for the 20 amino acids was treated as a group (PSSM_AA). The best model trained on the entire training dataset with parameters ntree = 80 and mtry = 7 was used to analyze the feature importance.

### Data and model availability

All PDB files including the HADDOCK-refined models, and PSSM files used in this work are available from the SBGrid Data repository^36^ (doi:10.15785/SBGRID/520). The iSEE predictor and features used for training and test are freely available on GitHub from https://github.com/haddocking/iSee.

## Results

### Training and validation of iSEE on a large diverse single point mutation dataset

We trained iSEE on a relatively large and diverse dataset consisting of experimental ΔΔG values for 1102 single point mutations in the interface of 57 dimer complexes. Among those, 656 mutations are in loops, 767 are non-ALA mutations, 376 correspond to small to large substitutions and 590 from large to small size. For each mutation, we extracted 31 energetics and conservation features (see Methods). A random forest (RF) model was trained and evaluated using 10-fold cross-validation(CV). The data are randomly divided into 10 parts, 9 of which are used for training and the left-out one for evaluating the performance of the trained RF model. This process was repeated 10 times to reduce the randomness of the data partition. From this training, a RF model with 80 trees and 7 randomly selected variables for each node achieved the best average root mean square error (RMSE) value **(Figure S1** in Supporting Information). The resulting best performing ΔΔG predictor, called iSEE, was compared with state-of-the-art ΔΔG predictors (see below).

iSEE’s prediction performance shows an average RMSE of 1.41±0.14 kcal mol^-1^ and a Pearson’s correlation coefficient (PCC) of 0.80±0.06 over the cross-validated sets **(Figure 2A)**. The predictor performs as well for ALA and non-ALA mutations, mutations inside and outside loops, and mutations corresponding to different changes in side-chain sizes (**Figure 2B-2D**). This indicates that our approach is not very sensitive to possible conformational changes coming from loop flexibility and is robust for different types of mutations. We further evaluated the applicability of iSEE to different types of protein complexes. Our results show that iSEE has a strong generalizability for predicting ΔΔG trends for mutations within complexes (**Figure S2 and S3**).

**Figure 2.**
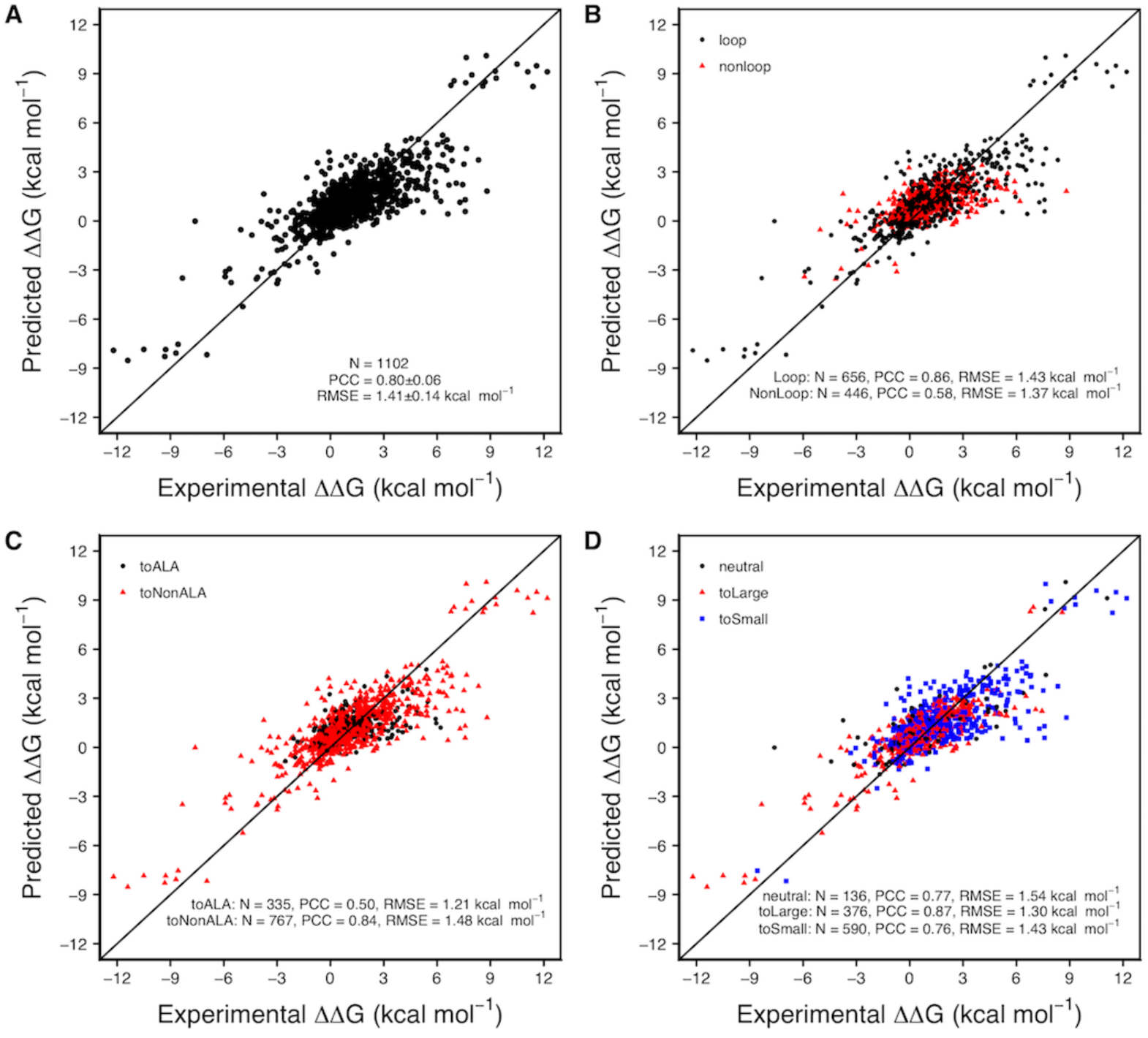
Correlations between predicted and experimental ΔΔG values for the training dataset consisting of 1102 single point mutations from the SKEMPI^14^/DACUM^27^ database. Ten times 10-fold cross-validation (CV) was applied during training, and the average of the CV predicted ΔΔG values are shown here for all mutations (A) and mutations classified as loop or non-loop (B), type of mutated amino acid (C), and change in amino acid size (D). The diagonal indicates an ideal prediction. PCC is the Pearson’s correlation coefficient and RMSE represents root mean squared error.

### iSEE competes with state-of-the-art ΔΔG predictors

We evaluated the performance of our iSEE ΔΔG predictor on the blind Benedix *et al.* NM dataset^8^ (see Methods) and compared it to several other state-of-the-art ΔΔG predictors based on empirical potentials or machine learning methods, which have been tested by Li et al.^9^ on the same data set. We only selected data from the NM data set for mutations that were not represented in the training data, which left 19 single point mutations for one complex (heterodimer, PDB ID: **1IAR**).

iSEE was compared to the following predictors:

- FoldX, which models free energy as a linear combination of multiple energy terms with weights optimized on a set of experimental ΔΔG values^6^.
- ZeMu, which can model conformational changes upon mutation using molecular dynamics simulations but relies on FoldX to predict ΔΔG^7^.
- CC/PBSA^8^, pred1^9^ and pred2^9^, which generate an ensemble of structures and apply a Molecular Mechanics - Poisson-Boltzmann Surface Area (MM-PBSA) approach to calculate the binding free energy.
- BeAtMuSiC, which is based on a linear combination of coarse grained statistical potentials^10^.
- mCSM^11^, a machine learning based approach, using distance-specific atom-contacts (calculated from the wild-type structures only) and pharmacophore changes of the mutation site as features of Gaussian processes to predict ΔΔG.
- BindProfX^19^, which is mainly based on evolutionary interface profile constructed from structural homologs, and combines the interface profile score with FoldX through a simple linear function to predict ΔΔG.

iSEE compares favorably with the eight other predictors considered here over the independent NM test set with a RMSE of 1.37 kcal mol^-1^ and a PCC of 0.73 (**Figure 3**), belonging to the top four predictors with PCCs over 0.70: BindProfX (0.81), iSEE (0.73), FoldX (0.72) and ZeMu (0.70). The ΔΔG predictions of the top four predictors are statistically significant with p values lower than 0.001, while other predictors have larger p values ranging from 0.006 to 0.523. Note that since CC/PBSA did use the NM data for training^8^, its performance might be over-estimated.

**Figure 3.**
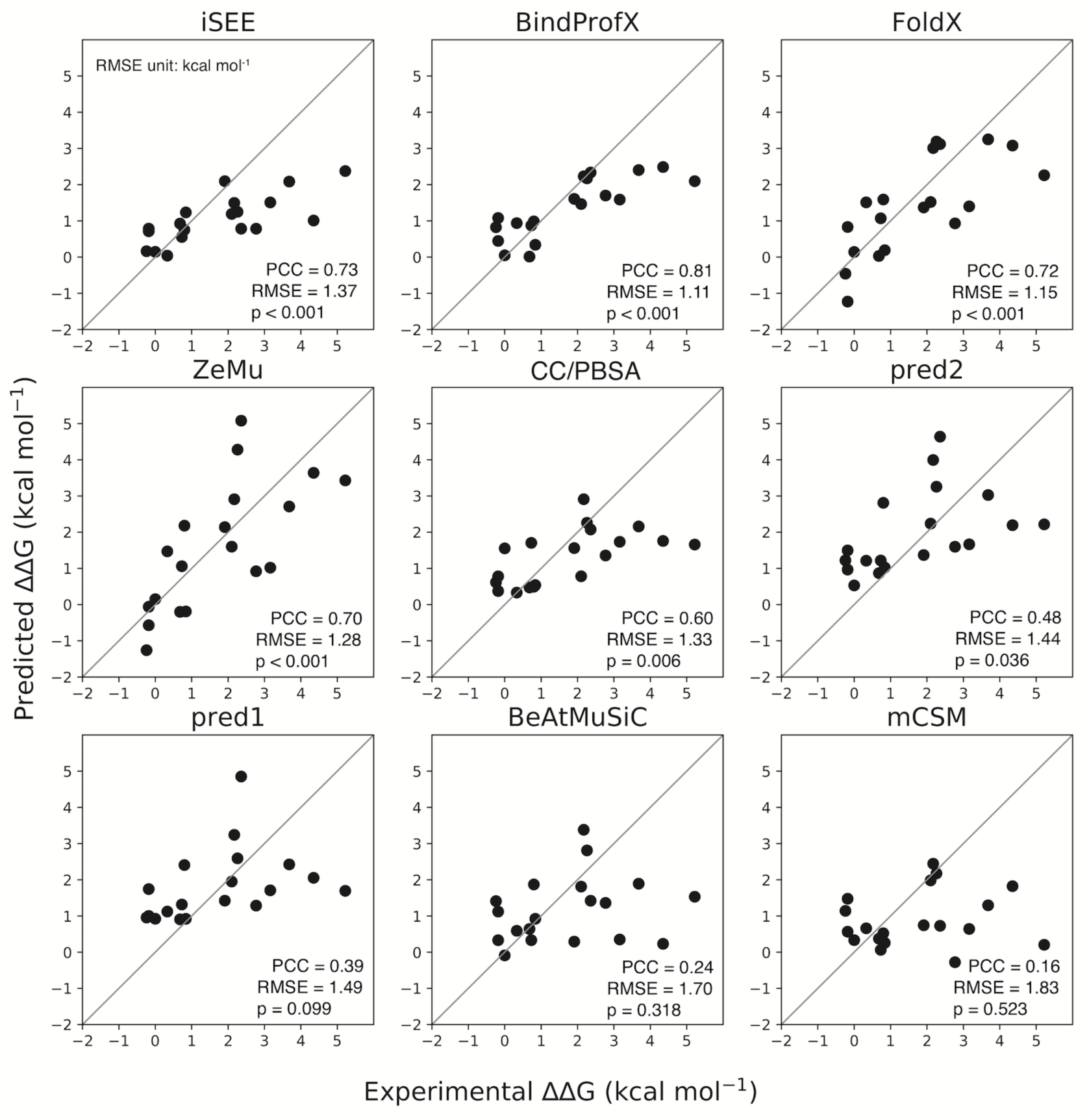
Predicted versus experimental ΔΔG for various ΔΔG predictors tested on a subset of the Benedix *et al.* NM dataset^8^ consisting of 19 mutations for one complex, non-overlapping with our training set. This subset was not used in any of the predictors, except for CC/PBSA. PCC is the Pearson’s correlation coefficient, p is two tailed p value of PCC, and RMSE represents root mean squared error.

### There is still plenty of room to further improve ΔΔG predictors

We benchmarked iSEE and three other ΔΔG predictors (FoldX, mCSM and BindProfX) on a much larger blind test dataset (the S540 dataset) constructed from the recently released SKEMPI 2.0 update. This dataset contains 540 mutations in 58 protein complexes that have not been seen by any predictor tested here. None of the four ΔΔG predictors performs well on this large blind test set (**Figure 4**). BindProfX performs best with an RMSE of 1.24 kcal mol^-1^ and PCC of 0.38 while iSEE achieves an RMSE of 1.32 kcal mol^-1^ and PCC of 0.27.

**Figure 4.**
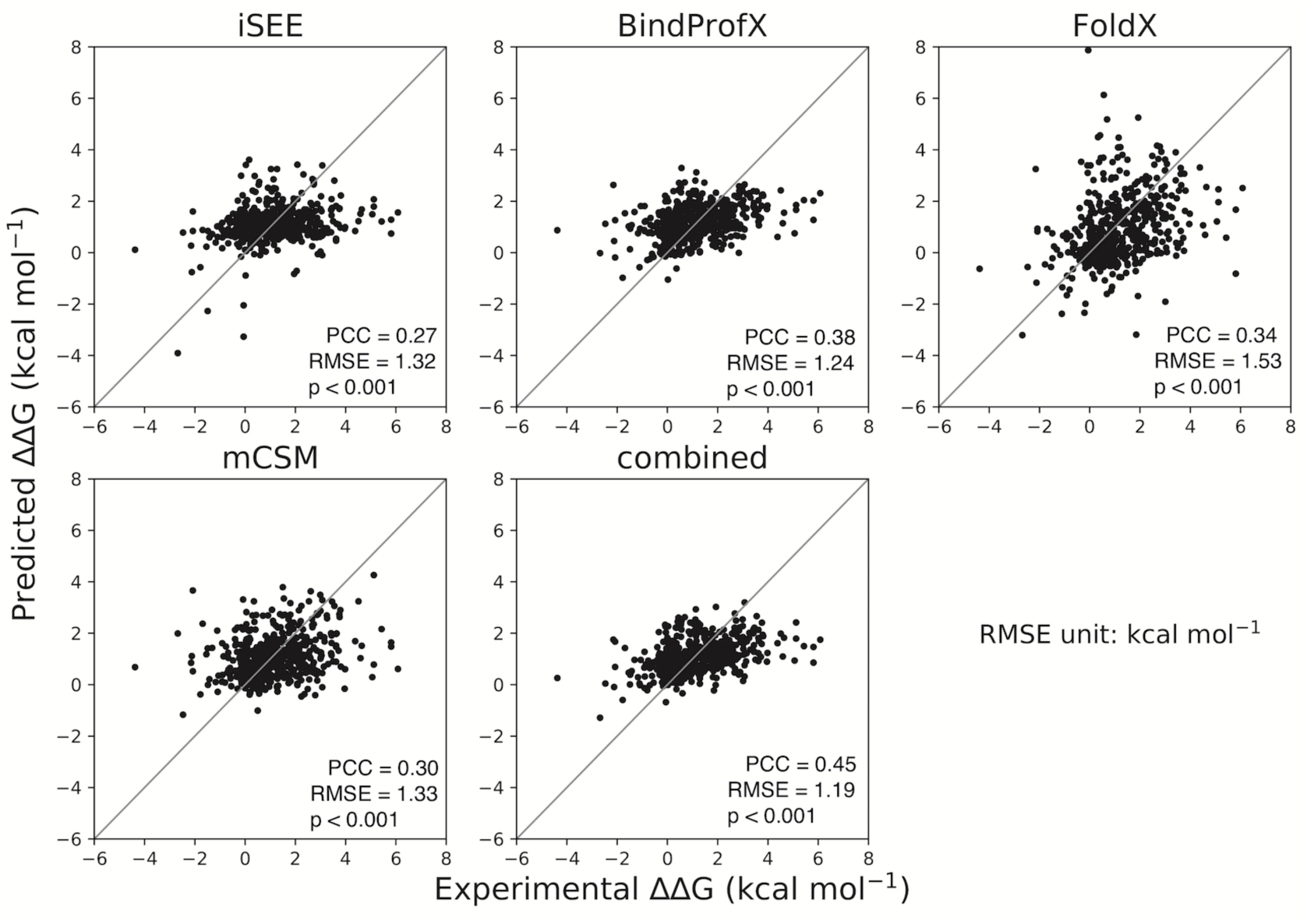
Correlations between predicted and experimental ΔΔG for various ΔΔG predictors tested on 540 mutations of SKEMPI 2.0. PCC is the Pearson’s correlation coefficient, p is two tailed p value of PCC, and RMSE represents root mean squared error.

To see if a combination of those ΔΔG predictors could improve the prediction performance, we simply averaged their predictions. The resulting combined predictor outperforms all the individual predictors with improved RMSE (1.19 kcal mol^-1^) and PCC (0.45) (**Figure 4**).

### Feature importance

We analyzed the importance of iSEE features for the prediction performance. This was done by calculating the averaged decrease of mean squared error for splitting on a given feature over all trees in the random forest model (see Methods). The results (**Figure 5**) reveal that the PSSM value of the wildtype amino acid (PSSM_wt) and the difference of PSSM values between mutant and wildtype residues (PSSM_diff) are the two most important features. They capture the evolutionary conservation of a specific amino acid at the mutation position and its change after mutation, respectively. PSSM has been proven to provide crucial information in various related topics, such as binding site predictions^37^ and hot-spot predictions^38^. The alignment depth does not seem to have much impact on the prediction performance (**Figure S4**). However, with most entries having over 300 sequences in their alignment a more systematic study should be performed to come to clear conclusions on this. The next most important feature is an energetic term, namely the change in intermolecular electrostatic energy calculated by HADDOCK between the mutant and wildtype complexes (Eelec_diff), followed by the PSSM information content (PSSM_IC). The latter captures the evolutionary conservation over all 20 types of amino acids that can potentially appear at the mutation position. The high importance of the PSSM_wt, PSSM_diff and PSSM_IC features indicate that evolutionary conservation is essential to quantitatively describe the effect of mutations on binding affinity.

**Figure 5.**
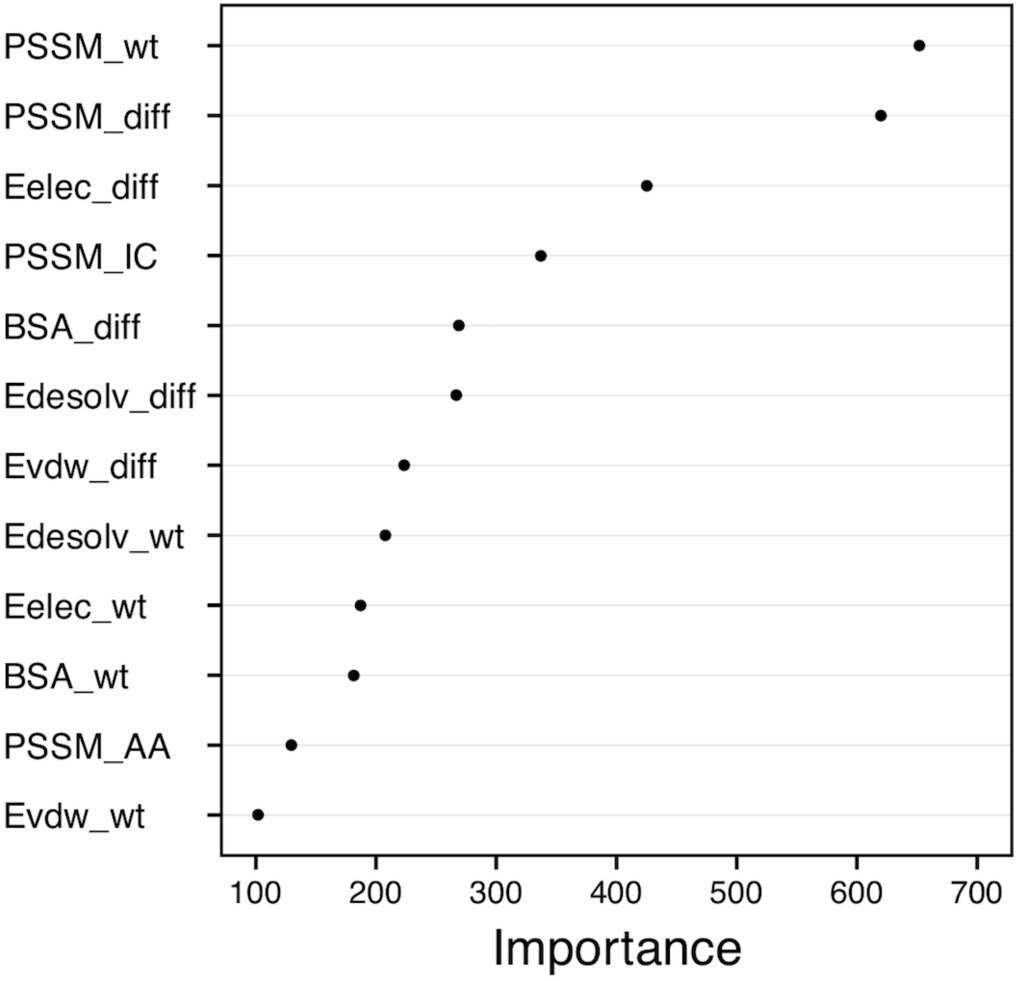
iSEE feature importance analysis. The importance value is measured as the decrease of mean squared prediction error when splitting on a given feature, averaged over all trees. The higher its value, the more important is the corresponding feature. The PSSM profile scores for the 20 amino acids are presented as a group in “PSSM_AA”.

## Discussion

We have developed a machine learning based ΔΔG predictor, iSEE, for quantitative prediction of the effects of single point mutations at the interface of a protein-protein complex. By combining structural, evolutionary and energetic features and training on a large and diverse dataset, our iSEE predictor not only demonstrated a consistent and high performance on various types of mutations during training, but also competed with state-of-the-art methods (based on empirical potential or machine learning models) on independent blind test datasets.

Compared with other machine learning methods, our predictor uses a rather small number of features, 31 in total which minimize the risk of overfitting (mCSM, for example, could use over 100 features). Evolutionary features, which benefit from the wealth of sequence data, are particularly sensitive to describe the impact of mutations on binding affinity as demonstrated by our feature importance analyses. The evolutionary conservation at both the amino acid type level (PSSM_wt and PSSM_diff) and mutation position level (PSSM_IC) were dominant amongst all iSEE features. Next to evolutionary features, energetic terms calculated with HADDOCK contribute to a quantitatively prediction of ΔΔGs, in particular the change of intermolecular electrostatic energy (Eelec_diff).

Unlike mCSM for which only wild-type structures are needed as input, iSEE does require the structures of both wildtype and mutant complexes. Models of the mutant complexes were obtained using the HADDOCK refinement server^28^. The robust prediction results for mutations in loop vs. non-loop and mutations with different residue size changes indicates that this approach—the short refinement in explicit solvent performed by HADDOCK—can handle a small degree of conformational changes and remove steric clashes. To explore whether using an ensemble of structural models instead of a single model would improve the prediction performance, we also trained and tested iSEE using the average features calculated from the top 4 models returned by the HADDOCK refinement server. iSEE seems rather robust with respect to small conformational differences that might affect the energetic terms since using values from the top-ranked model or averages over the best 4 does not have any significant impact on its performance (**Table S1**). More systematic analyses are however needed to draw solid conclusions on this point.

With the recent release of SKEMPI 2.0, it becomes possible to benchmark current ΔΔG predictors on a large and novel blind dataset. Our benchmarking results on a set of 540 mutations show that all state-of-the-art ΔΔG predictors do not perform well with PCCs lower than 0.4. This indicates there is still a plenty of room for further improvements. Interestingly, averaging the predictions from the different ΔΔG predictors generated an improved prediction performance, indicating the various ΔΔG predictors might use complementary features. This should be useful for further development and improvement of ΔΔG predictors.

## Acknowledgement

This work was supported by the European H2020 e-Infrastructure grant BioExcel (grant no. 675728). CG acknowledges financial support from the China Scholarship Council (grant no. 201406220132). AV acknowledges financial support from the European Union’s H2020 Marie Sklodowska-Curie Individual Fellowships (grant no. BAP-659025). LX acknowledges financial support from the Dutch Foundation for Scientific Research (Veni grant 722.014.005). We thank Dr. I.S. Moreira, P. Koukos and F. Ambrosetti for fruitful discussions and help about datasets and hot-spots. We also acknowledge the use of software from the SBGrid consortium^33^ for various analysis tasks. This work has no conflict of interest.

## Supporting material

### SI Results

#### Evaluation of iSEE’s applicability to different protein complexes

The original cross-validation described in the main text splits the dataset based on mutations and not complexes. A consequence of this is that related mutations could appear for a given complex in both the training and test sets, which might lead to an overestimation of the performance. To check for this, we selected 13 complexes from the training dataset with more than 20 mutations per complex. We then compared the prediction performance of iSEE on leave-one-complex-out cross-validation and leave-one-mutation-out cross-validation. For the leave-one-complex-out CV, all mutations that belong to a specific complex were hold out as independent test data, and the model was trained on mutations from all the remaining complexes. For the leave-one-mutation-out CV, each mutation in a complex was, in turn, hold out as a test data and the remaining mutations of that complex and the mutations of all other complexes were used for training.

The prediction performance on each complex between leave-one-complex-out CV and leave-one-mutation-out CV is compared in **Figure S2A**. Both cross-validation schemes show similar performances as clearly seen from the similar PCC values (with a p-value of 0.127 of Wilcoxon signed rank test). This indicates that iSEE has a strong generalizability on predicting the trend of ΔΔG values among a group of mutations. The Wilcoxon test applied on RMSE values gives, however, a p-value of 0.008, indicating a small significant difference in prediction error for each complex between the two cross-validation schemes.

We can also see that the PCC values are not uniform over various complexes, covering a large range from −0.01 to 0.97. iSEE shows varying prediction performance on different complexes. This observation is not correlated to the binding affinity of the wildtype complex (PCC = 0.09, data not shown). However, the low PCC values mainly resulted from the complexes that have a narrow range of ΔΔG values (≤|0.5| kcal mol^-1^) (**Figure S2B and S3**), which are close to the experimentally measured ΔΔG error of about 0.5 kcal mol^−1^[1].

**Figure S1.**
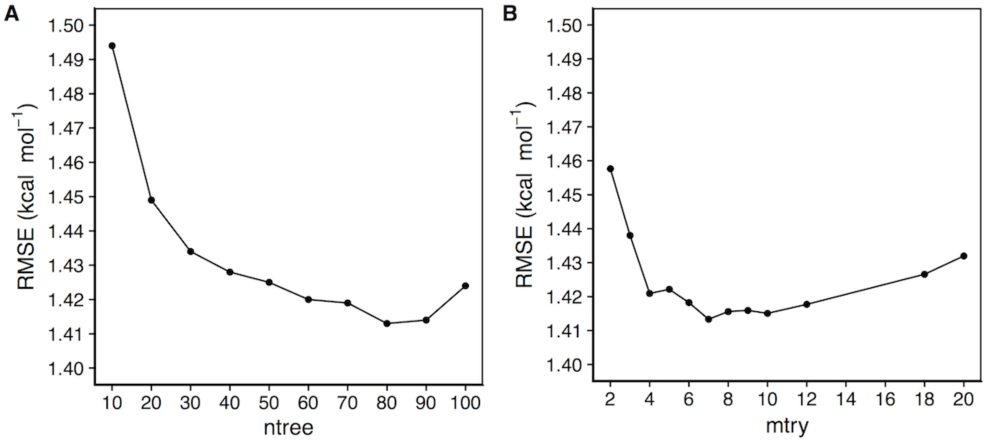
Cross-validation performance for different ntree and mtry settings. Given a specific ntree value, the random forest model is optimized on different mtry values by cross-validation on the training dataset. The best average RMSE for each ntree value is shown in (A). The lowest RMSE was obtained for ntree **=** 80. The RMSE values for different mtry values for ntree = 80 are shown in (B). The best RMSE was obtained for mtry = 7. The final iSEE predictor uses the parameters of ntree = 80 and mtry = 7.

**Figure S2.**
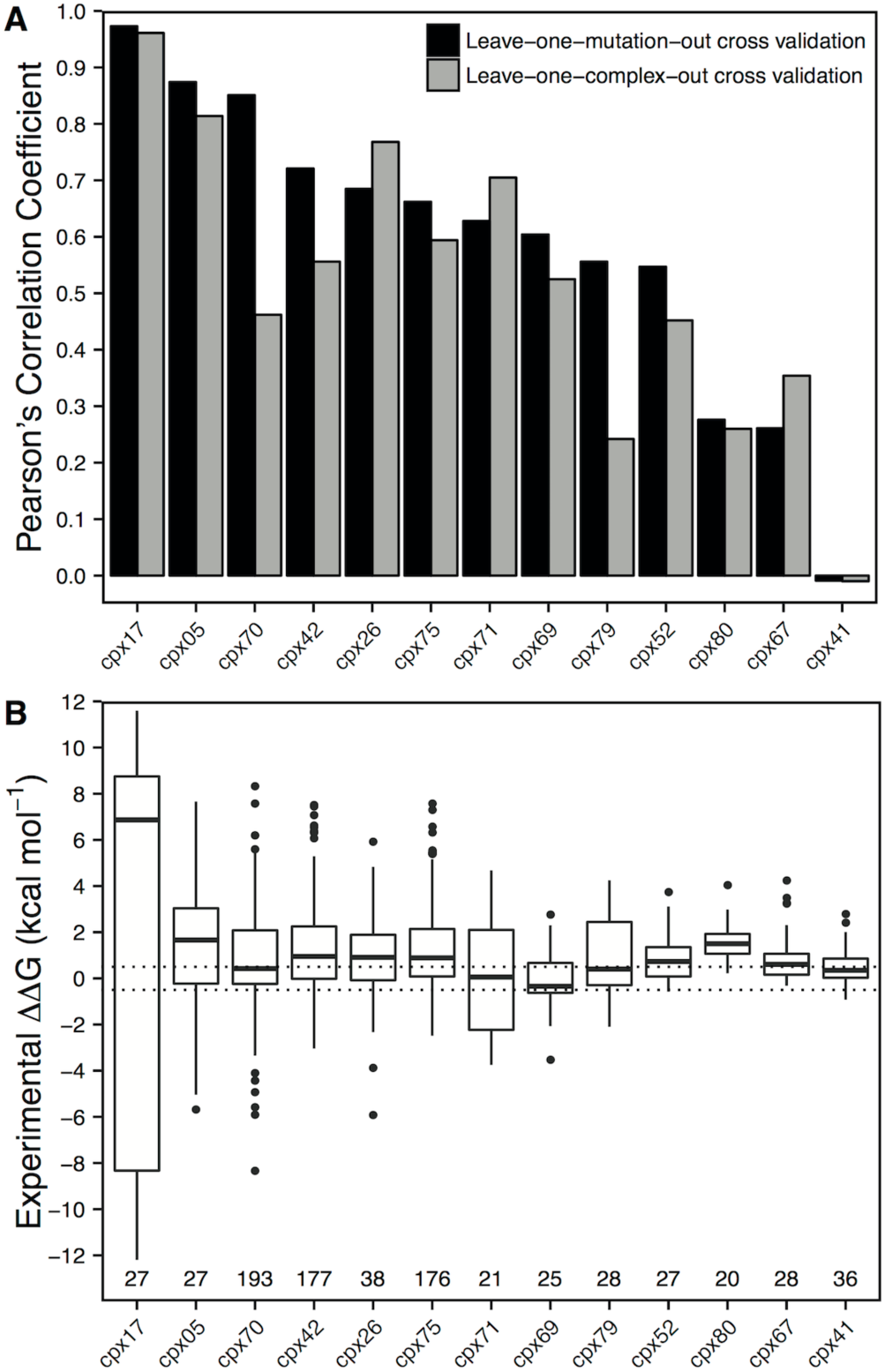
Evaluation of iSEE generalizability at the protein complex level. **(A)** Comparison of Pearson’s correlation coefficients for each complex between leave-one-mutation-out cross-validations and leave-one-complex-out cross-validations. **(B)** Distribution of experimental ΔΔG values and number of mutations for each complex. Dotted lines denote ΔΔG of +0.5 and −0.5 kcal mol^-1^. The complexes were ordered from left to right in order from high to low Pearson’s correlation coefficients of leave-one-mutation-out cross-validations.

**Figure S3.**
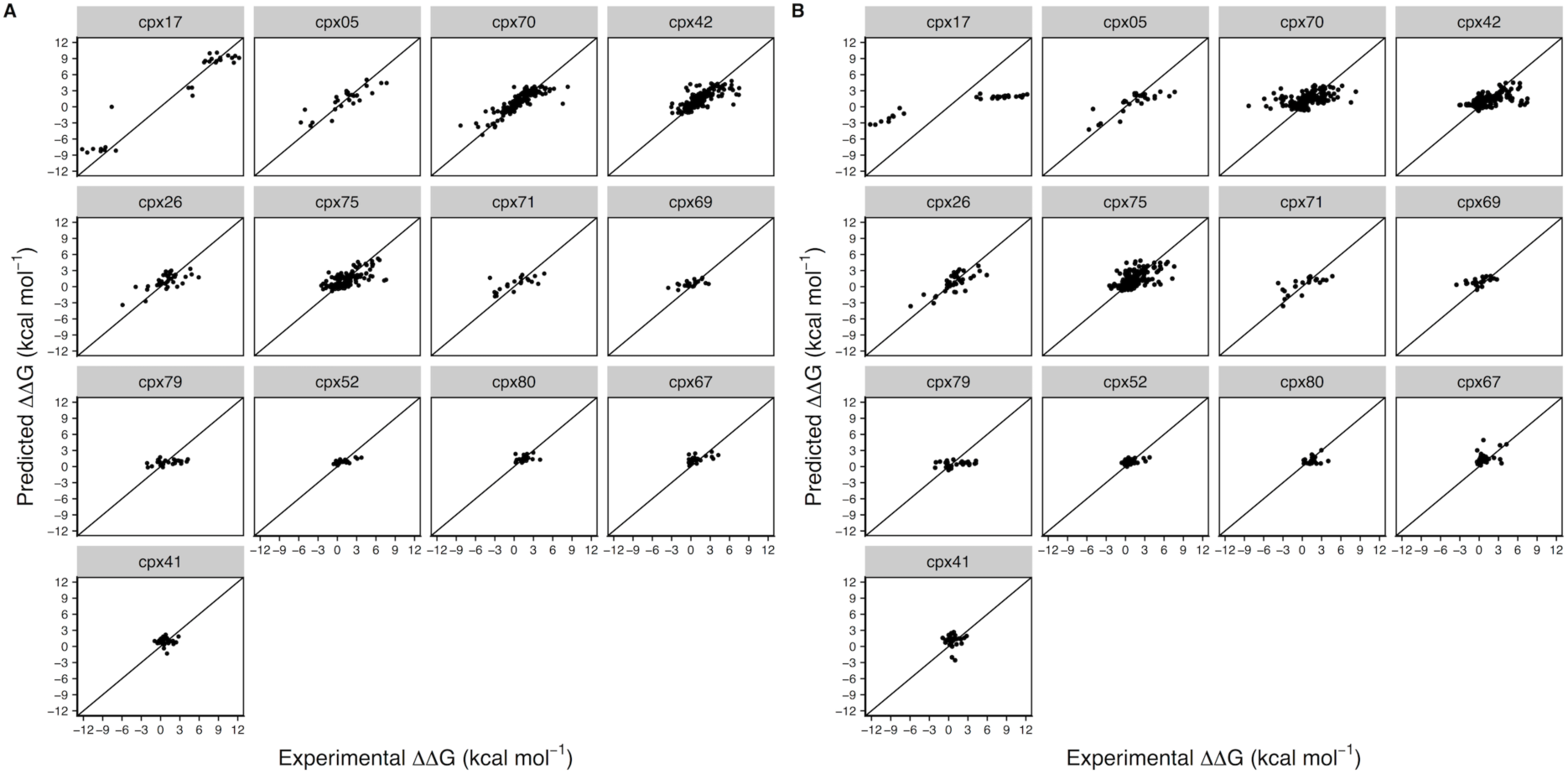
Correlations between predicted and experimental ΔΔG for each type of complex performing **(A)** leave-one-mutation-out and **(B)** leave-one-complex-out cross-validation. The complex types are ordered from left to right based on decreasing Pearson’s correlation coefficients of leave-one-mutation-out cross-validations.

**Figure S4.**
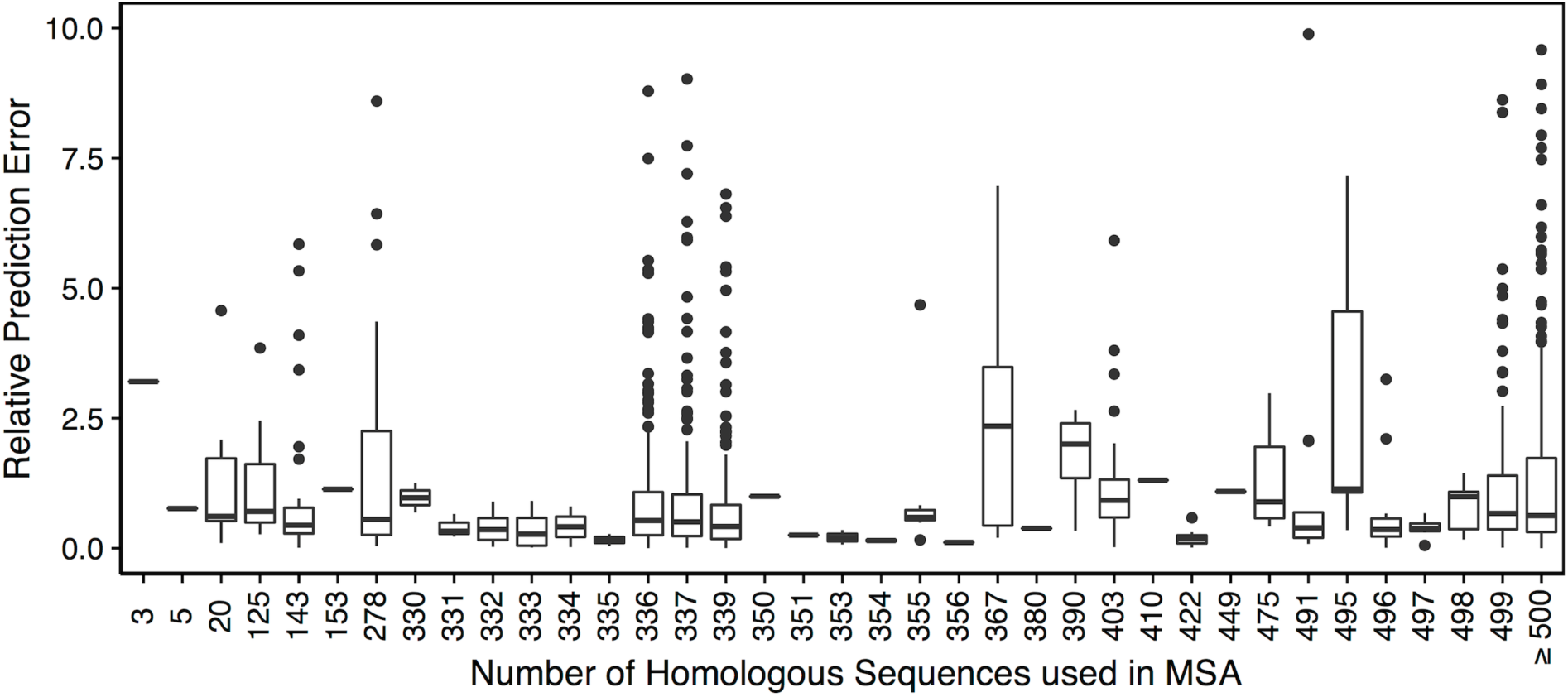
iSEE prediction’s performance as a function of the multiple sequence alignment depth used for constructing the PSSM. The performance is reported here as the relative prediction error: | (ΔΔGpred - ΔΔGexp) / ΔΔGexp | from cross-validations using the SKEMPI dataset. For clarity, the y axis scale excludes 25 mutations with relative prediction errors larger than 10. Further, 12 mutations with ΔΔGexp equal to zero are not shown here because we cannot calculate relative prediction errors for them.

**Table S1.** Performance of iSEE using different number of structural models from the HADDOCK refinement. The structure and energy features of iSEE were calculated using only the top (best ranked) structural model (iSEE_top1) and the average of features over the top four models (iSEE_top4).

|  | Method | RMSE<br>(kcal mol<sup>-1</sup>) | PCC |
| --- | --- | --- | --- |
| Training and Cross validation | iSEE_top1 | 1.41±0.14 | 0.80±0.06 |
|  | iSEE_top4 | 1.42±0.13 | 0.80±0.05 |
| Test on NM dataset | iSEE_top1 | 1.37 | 0.73 |
|  | iSEE_top4 | 1.20 | 0.77 |
| Test on S540 dataset | iSEE_top1 | 1.32 | 0.27 |
|  | iSEE_top4 | 1.39 | 0.23 |

**Table S2.** iSEE features.

| Feature Type | Feature Name |  | Comments |
| --- | --- | --- | --- |
| Structure | BSA_wt |  | Buried surface area of the wildtype complex |
|  | BSA_diff |  | = BSA_mut – BSA_wt |
| Energy | Evdw_wt |  | Intermolecular van de Waals energy of the wildtype complex |
|  | Eelec_wt |  | Intermolecular electrostatic energy of the wildtype complex |
|  | Edesolv_wt |  | Desolvation energy of the wildtype complex |
|  | Evdw_diff |  | = Evdw_mut – Evdw_wt |
|  | Eelec_diff |  | = Eelec_mut – Eelec_wt |
|  | Edesolv_diff |  | = Edesolv_mut – Edesolv_wt |
| Evolution | PSSM_wt |  | PSSM value for the wildtype amino acid type at the mutation position |
|  | PSSM_diff |  | = PSSM_mut – PSSM_wt |
|  | PSSM_IC |  | Information content at the mutation position |
|  | PSSM_AA | A, R, N, D,<br>C, Q, E, G,<br>H, I, L, K, M,<br>F, P, S, T,<br>W, Y, V | PSSM profile for 20 amino acid types at the mutation position |

**Table S3.**
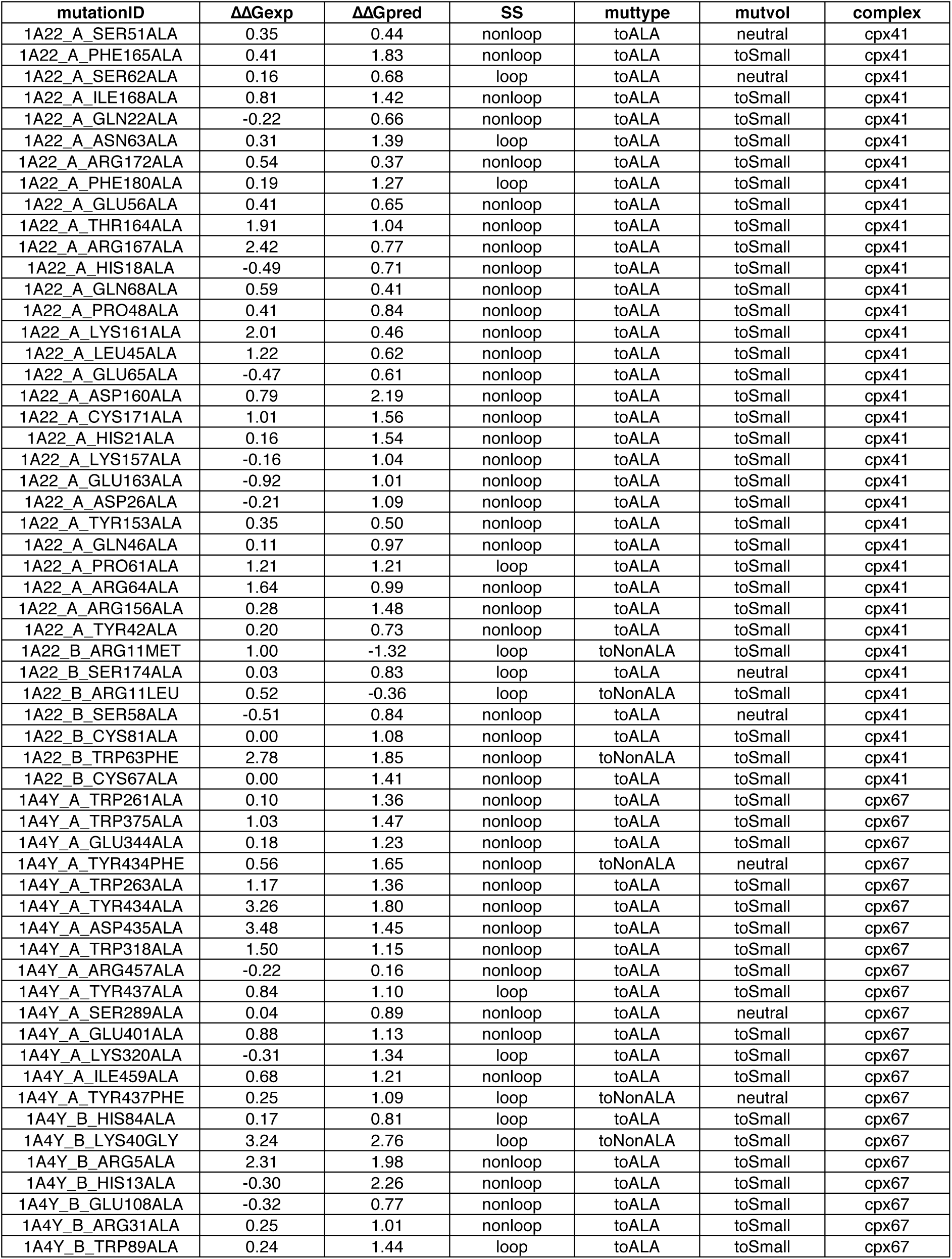
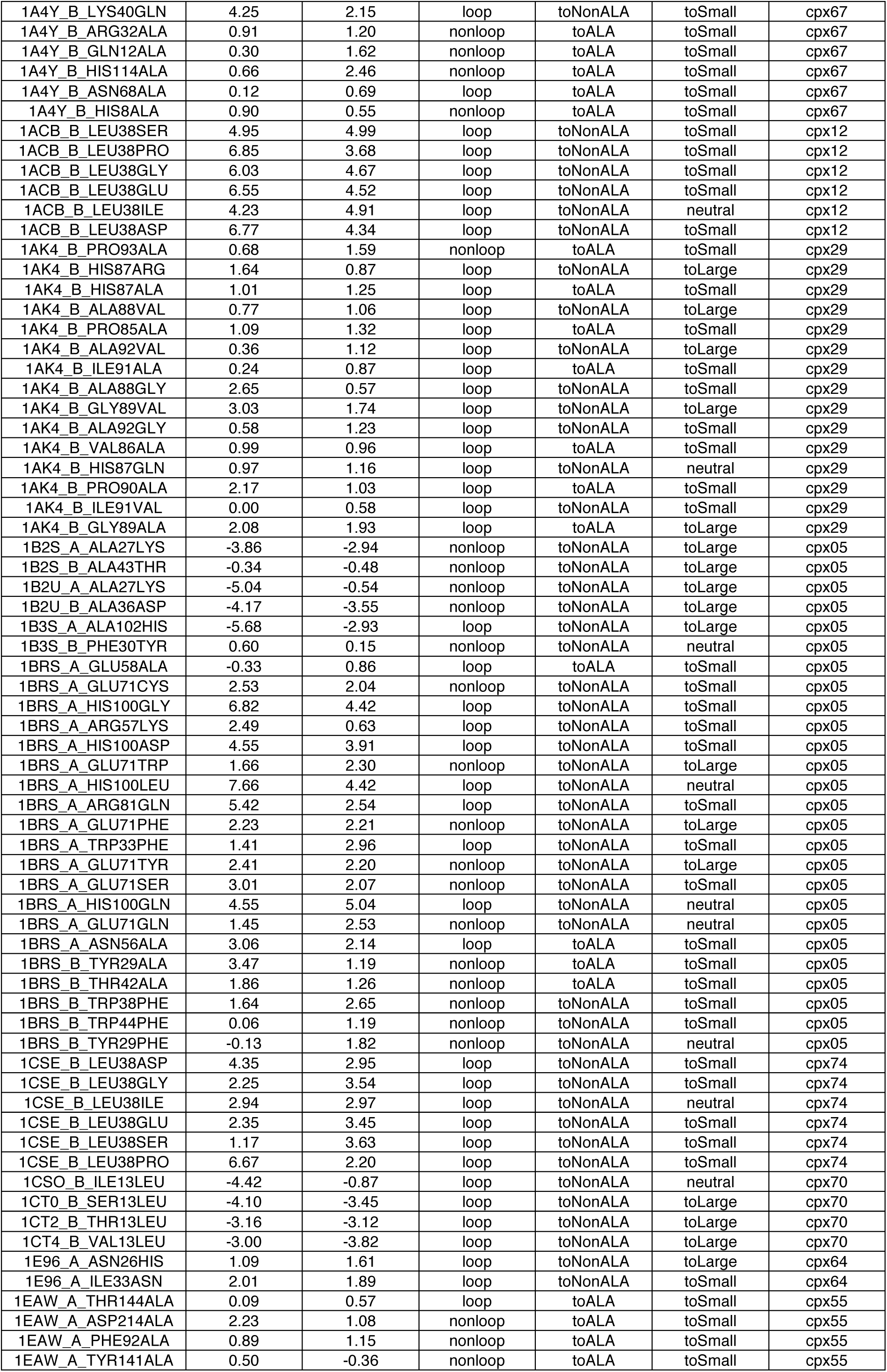
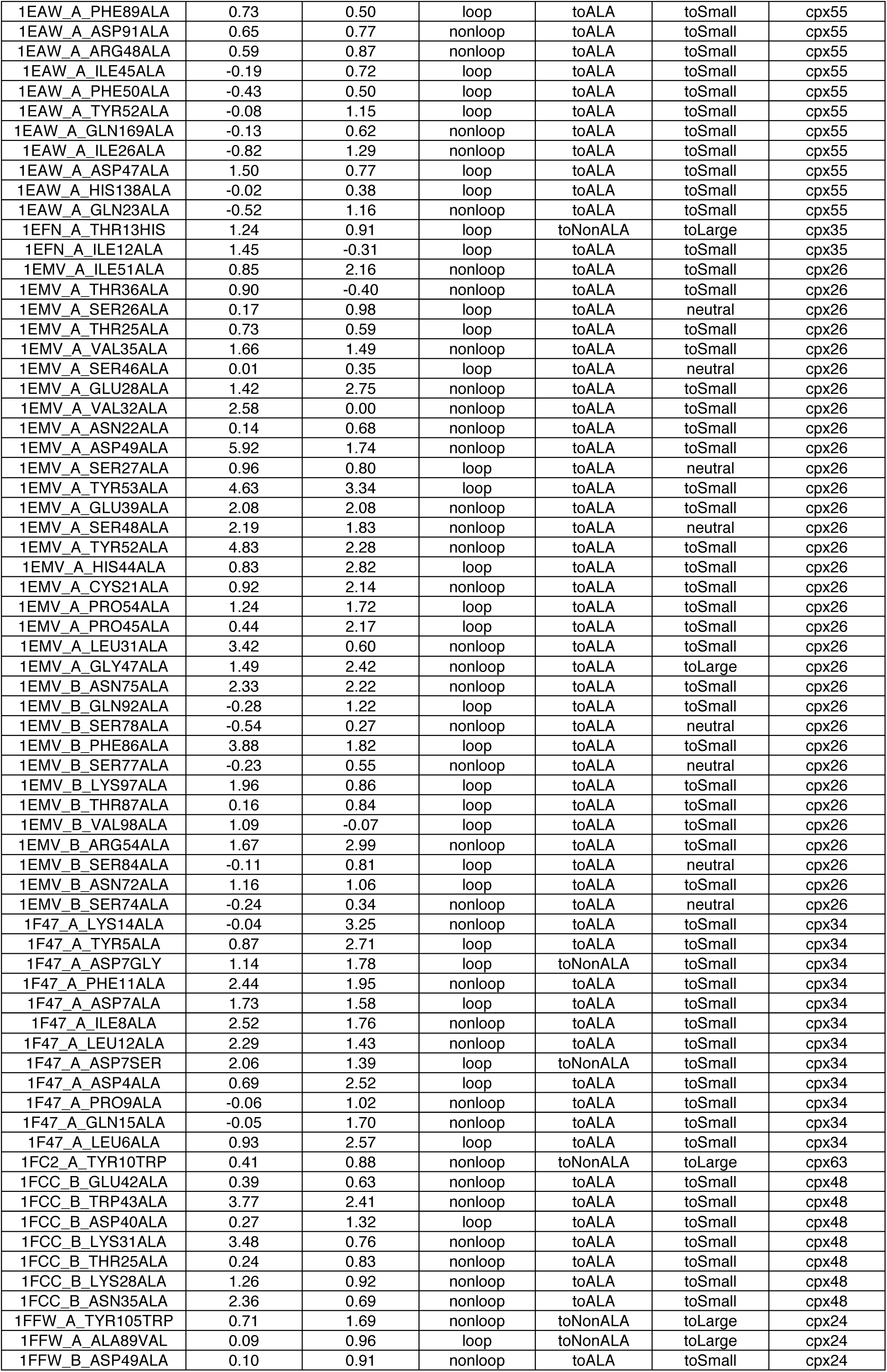
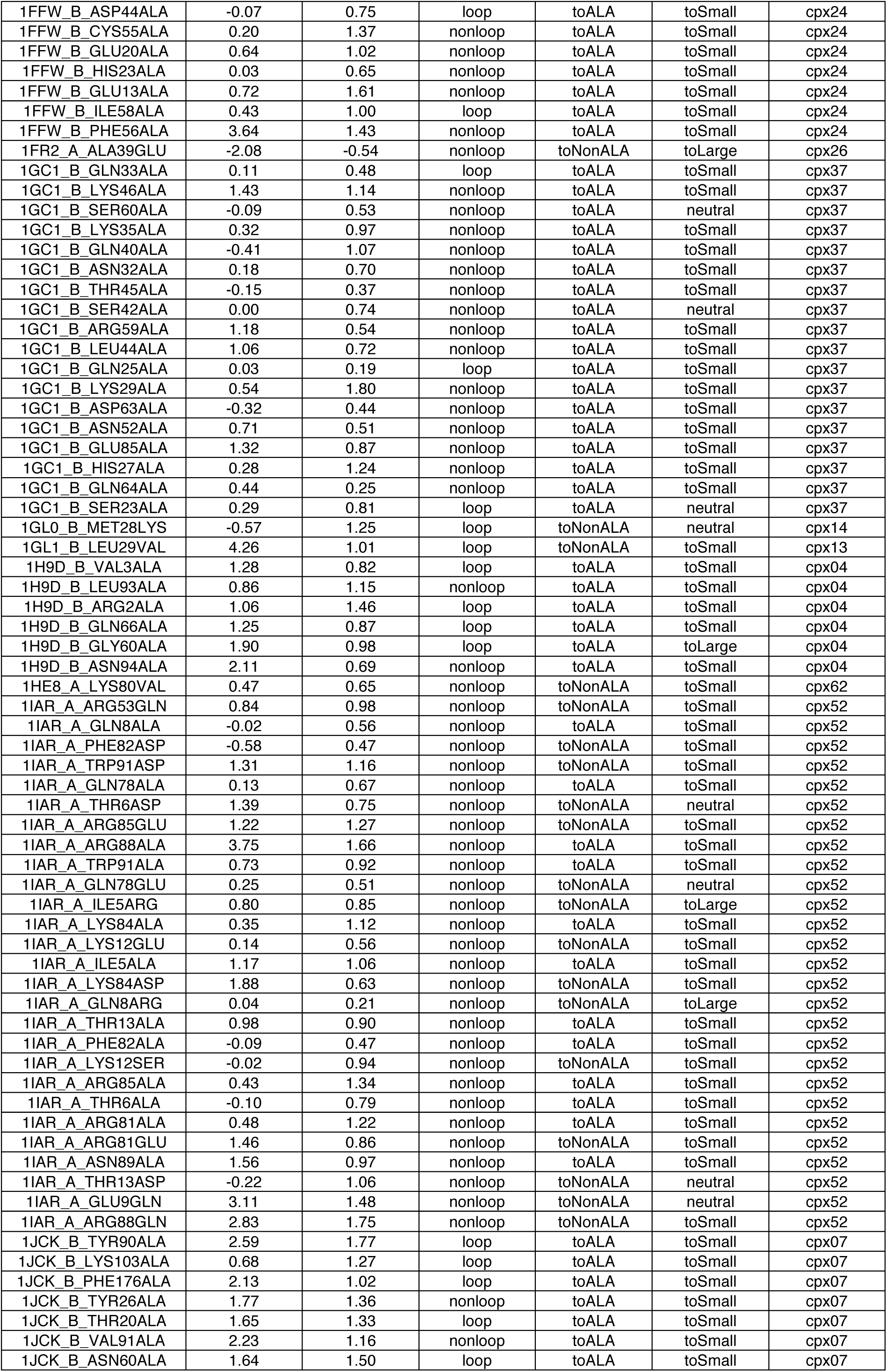
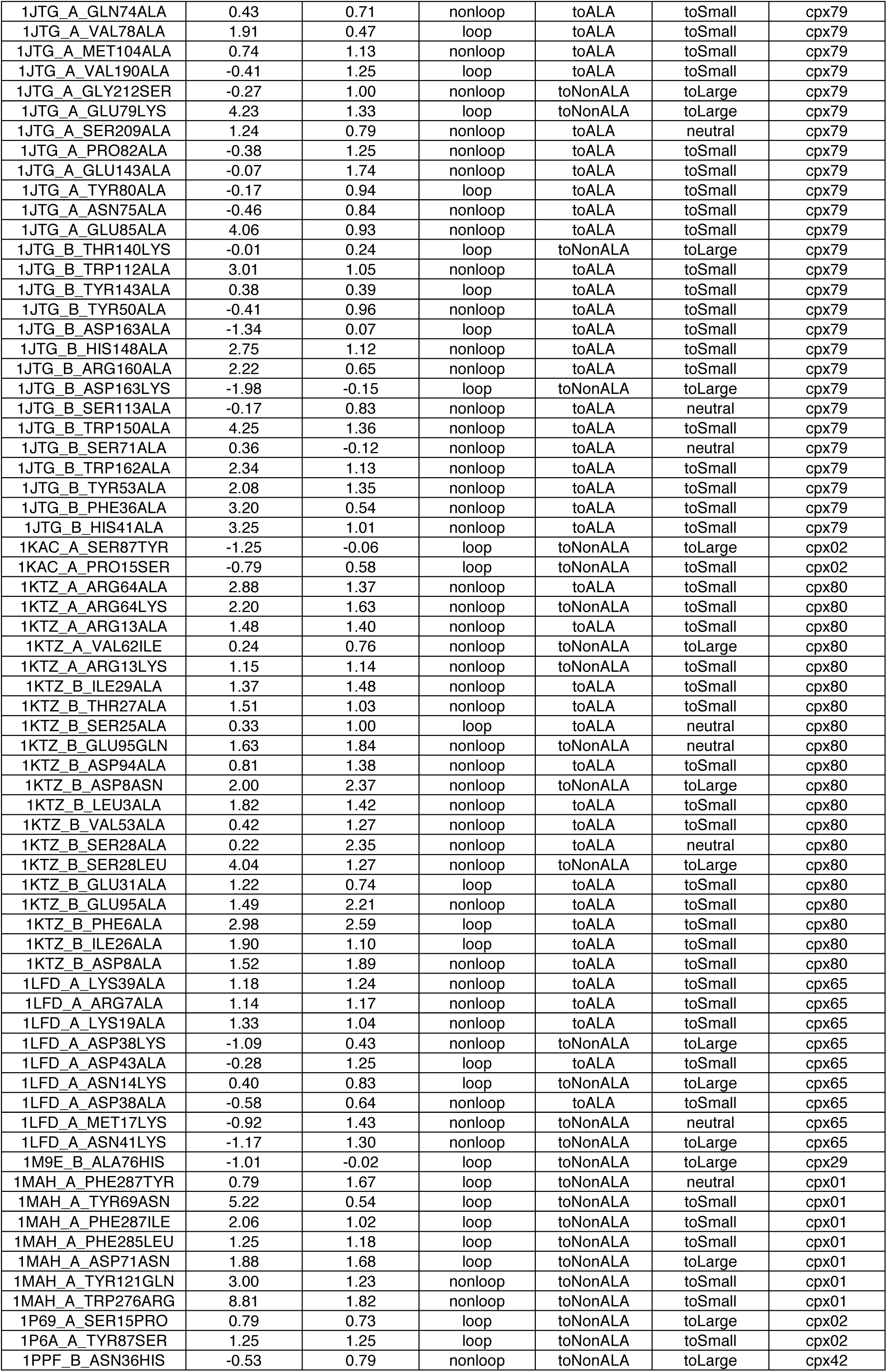
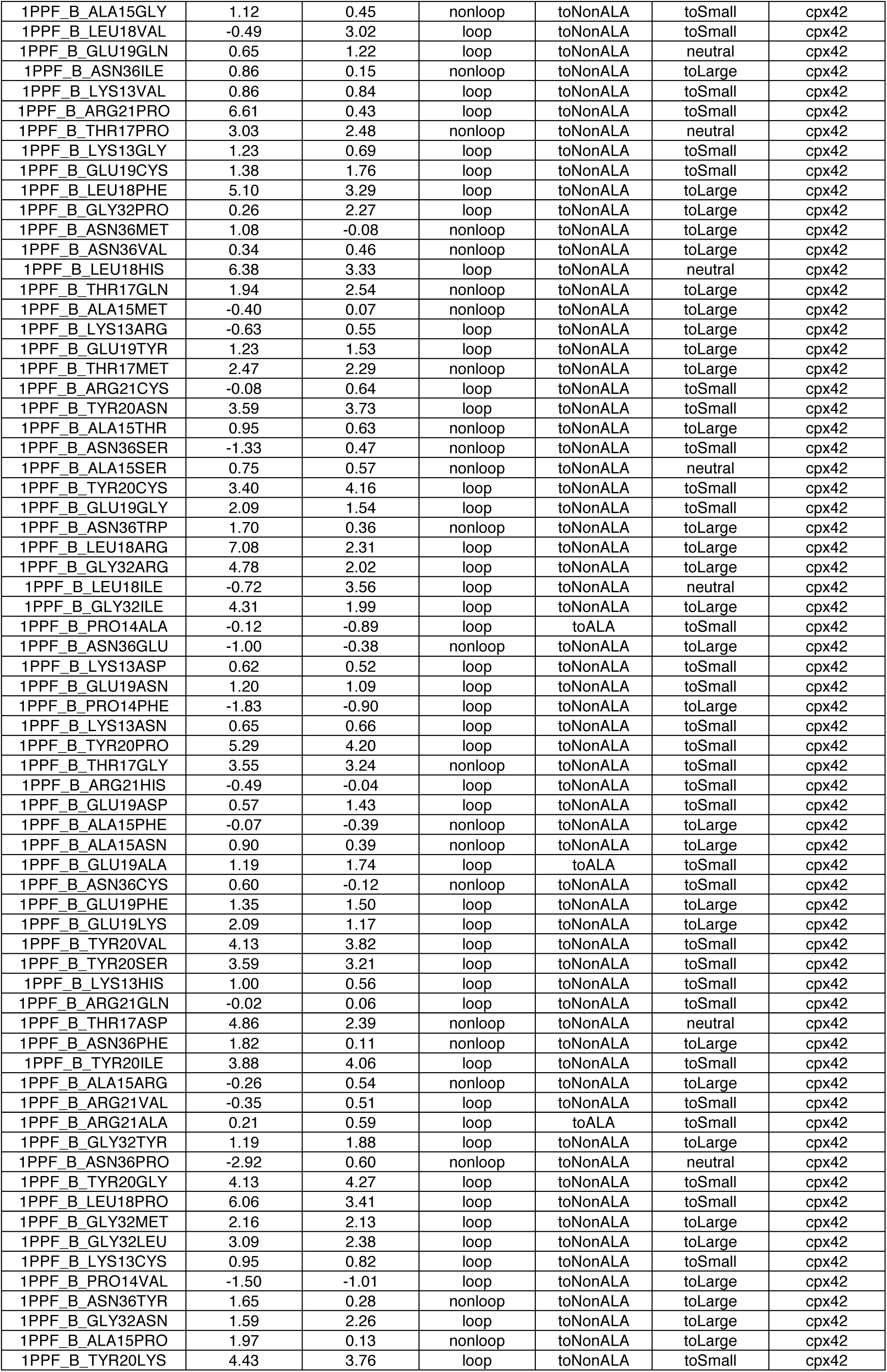
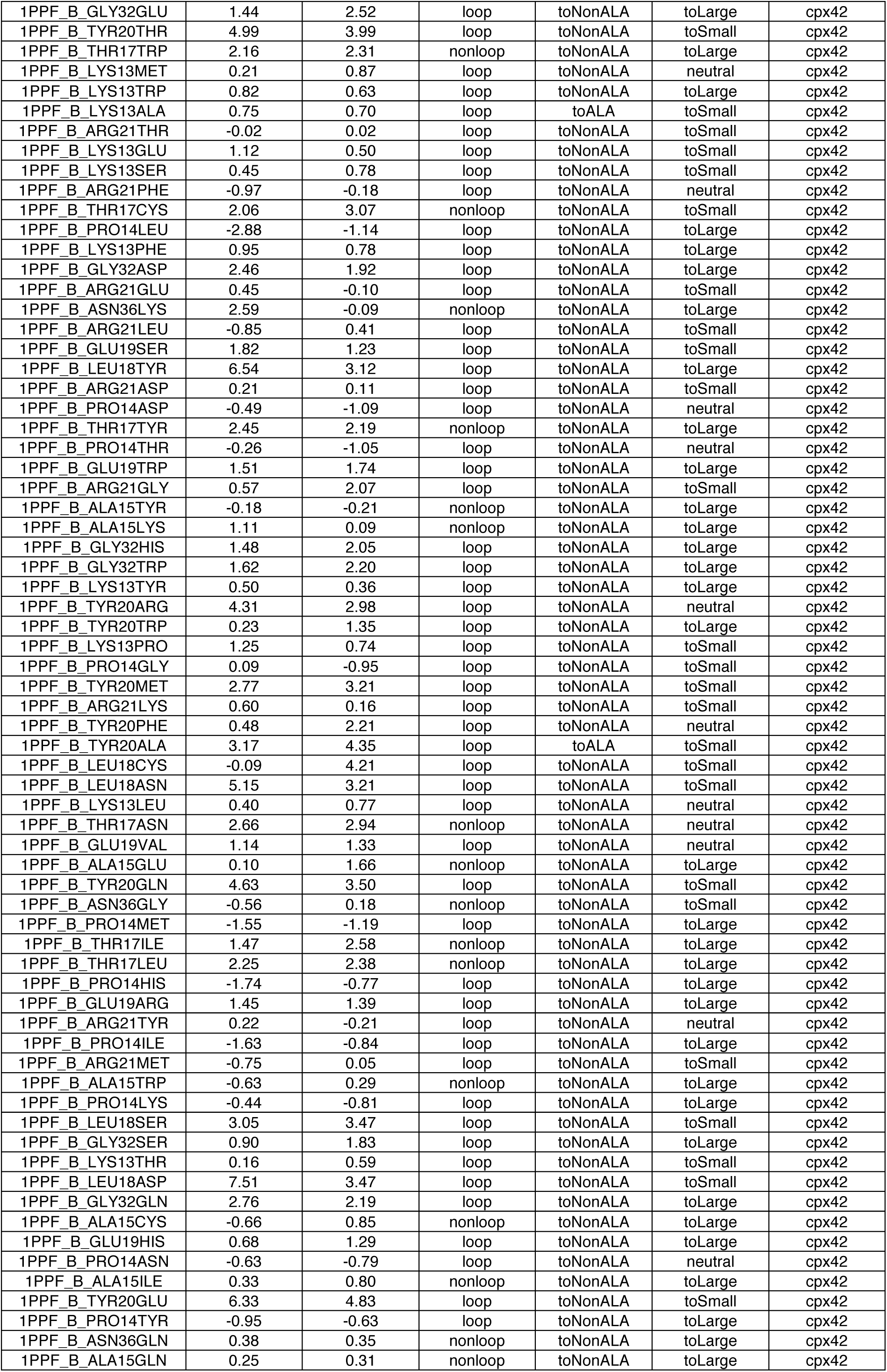
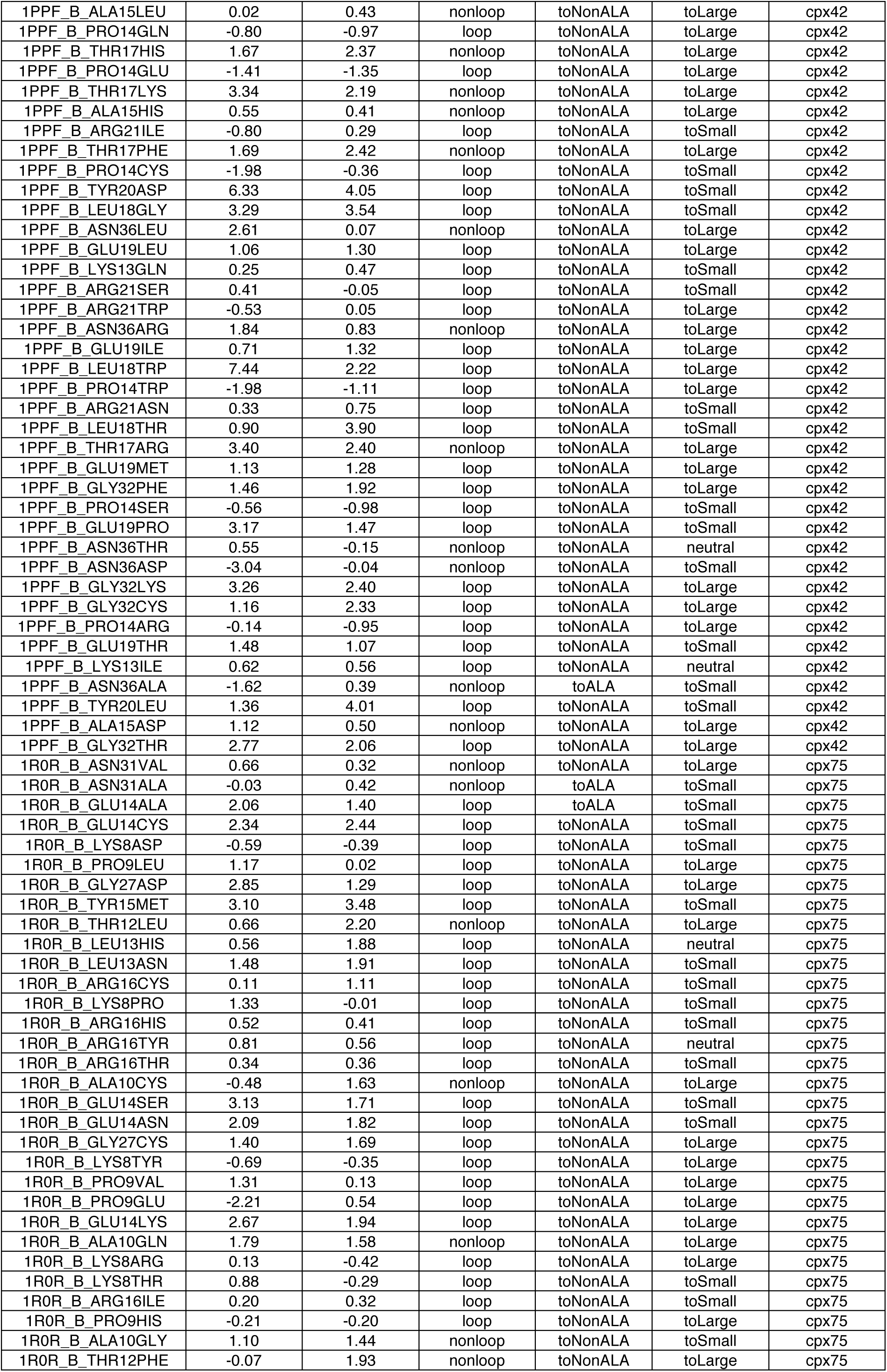
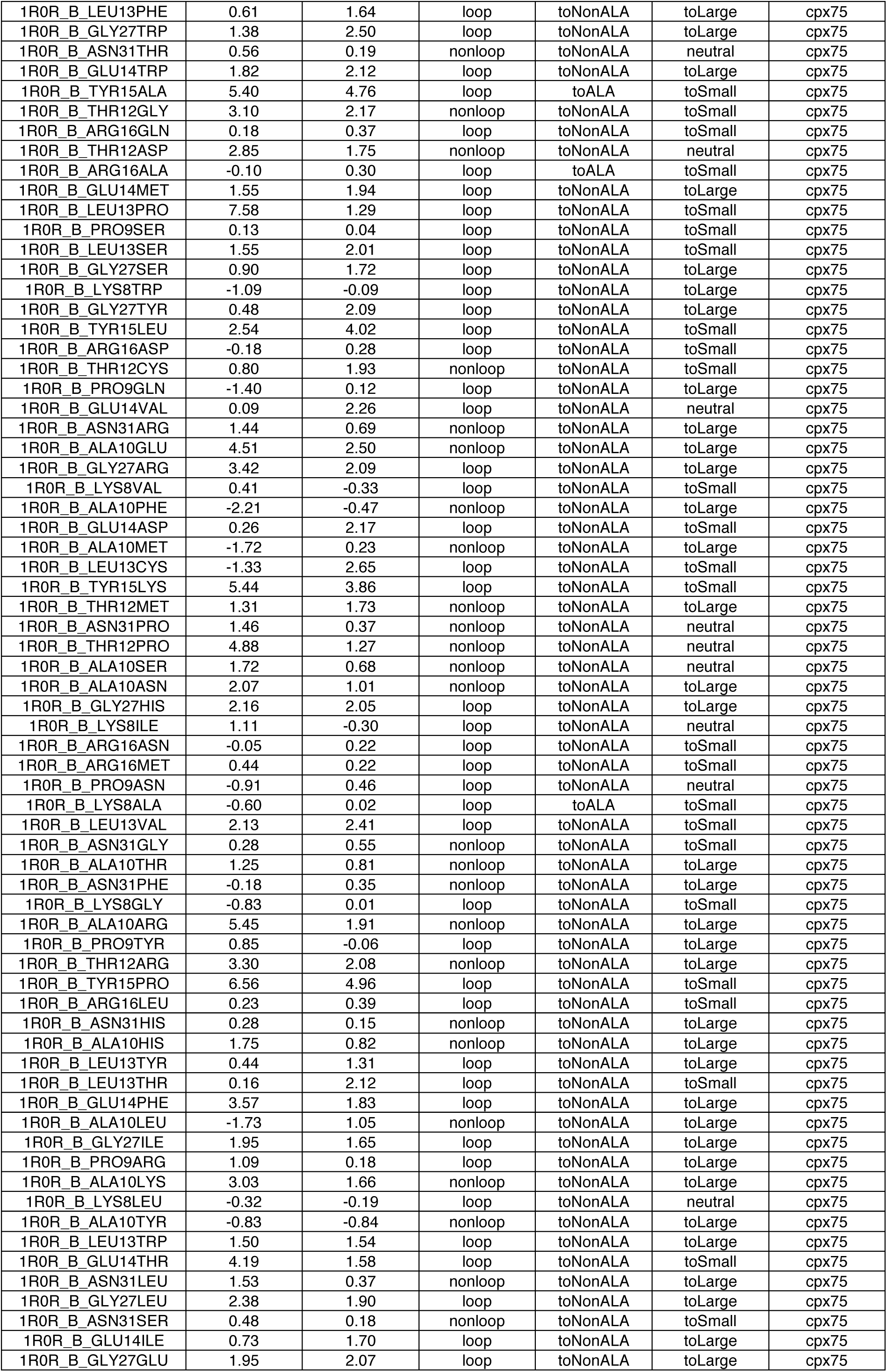
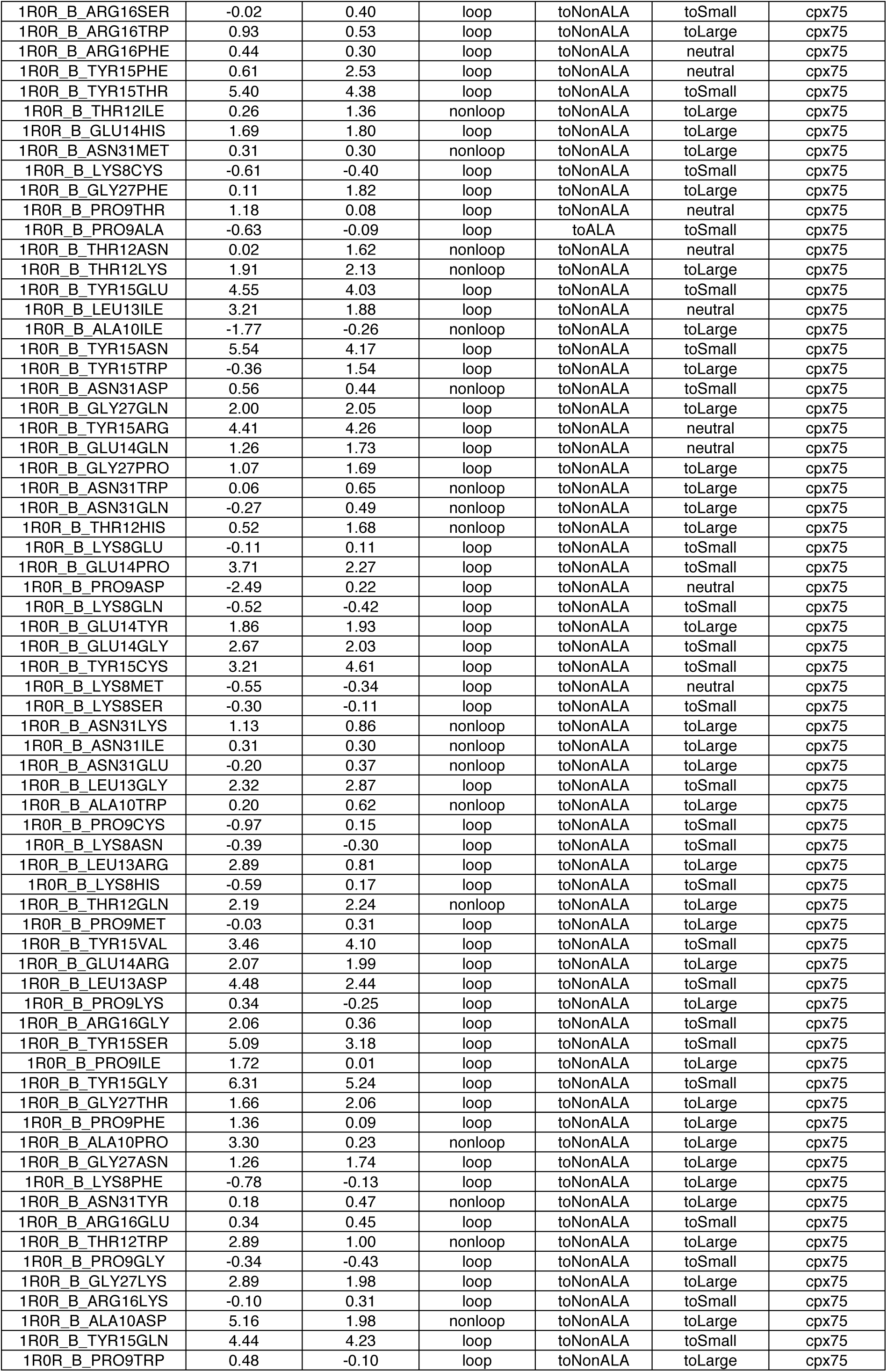
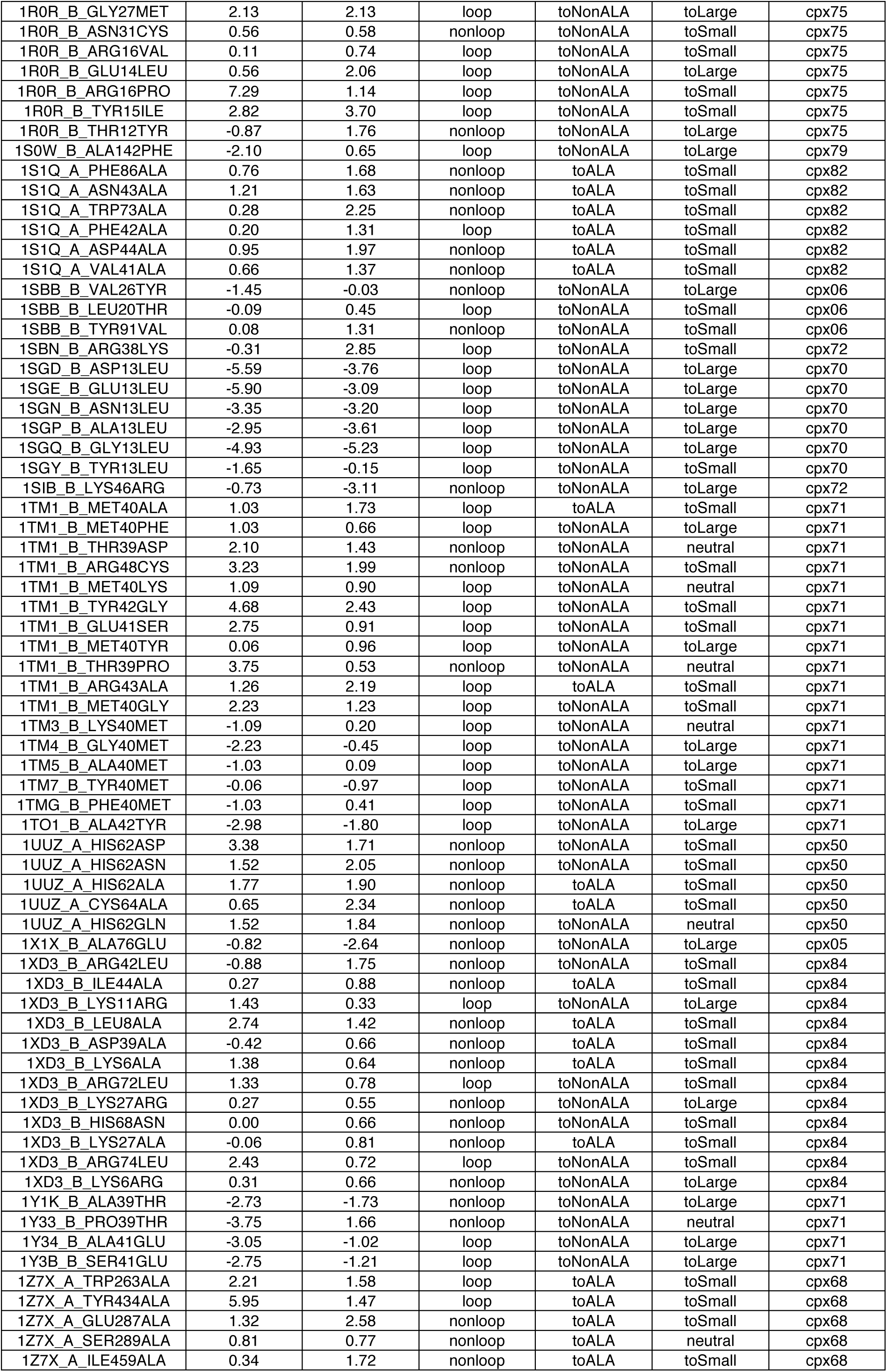
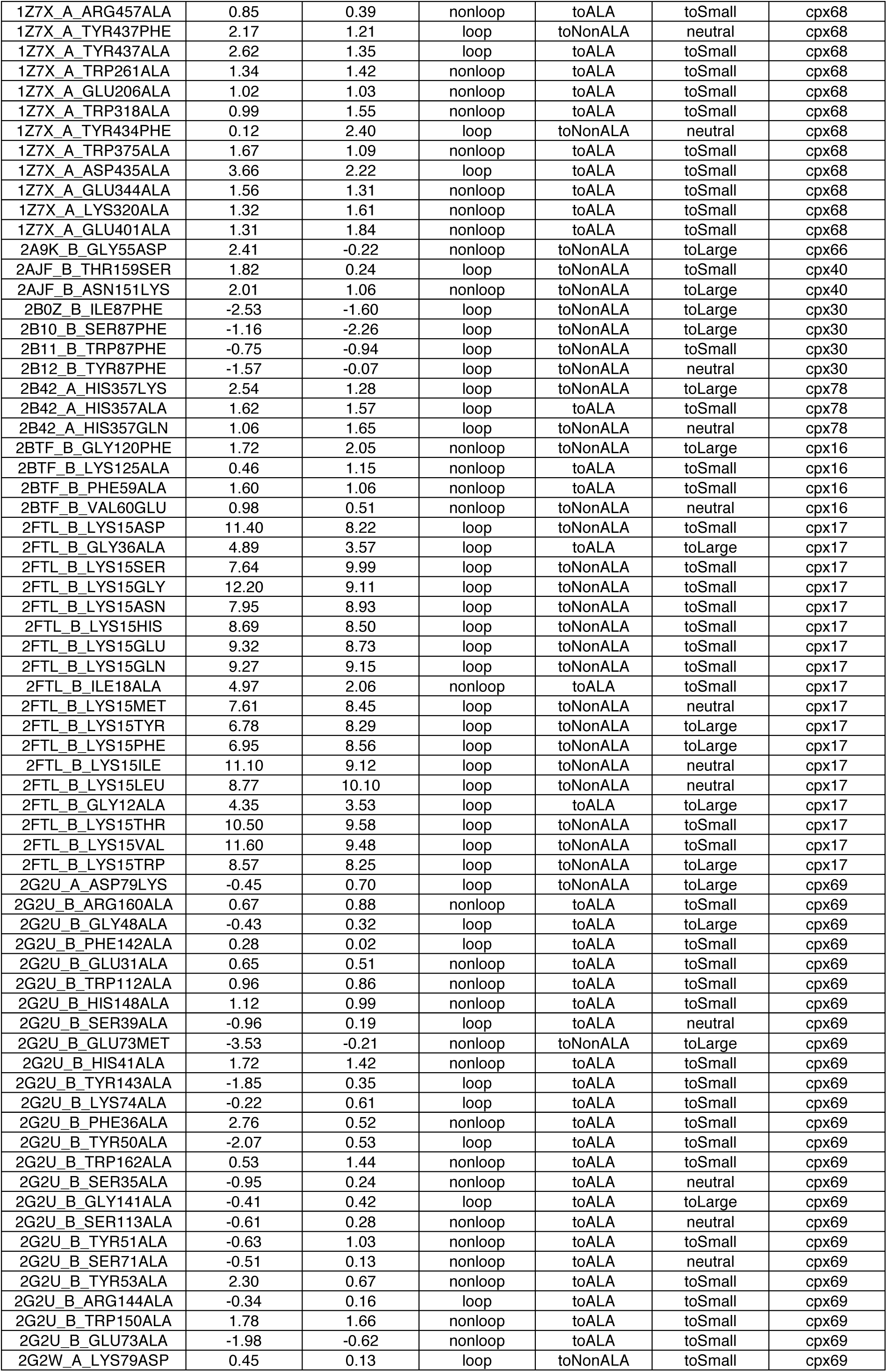
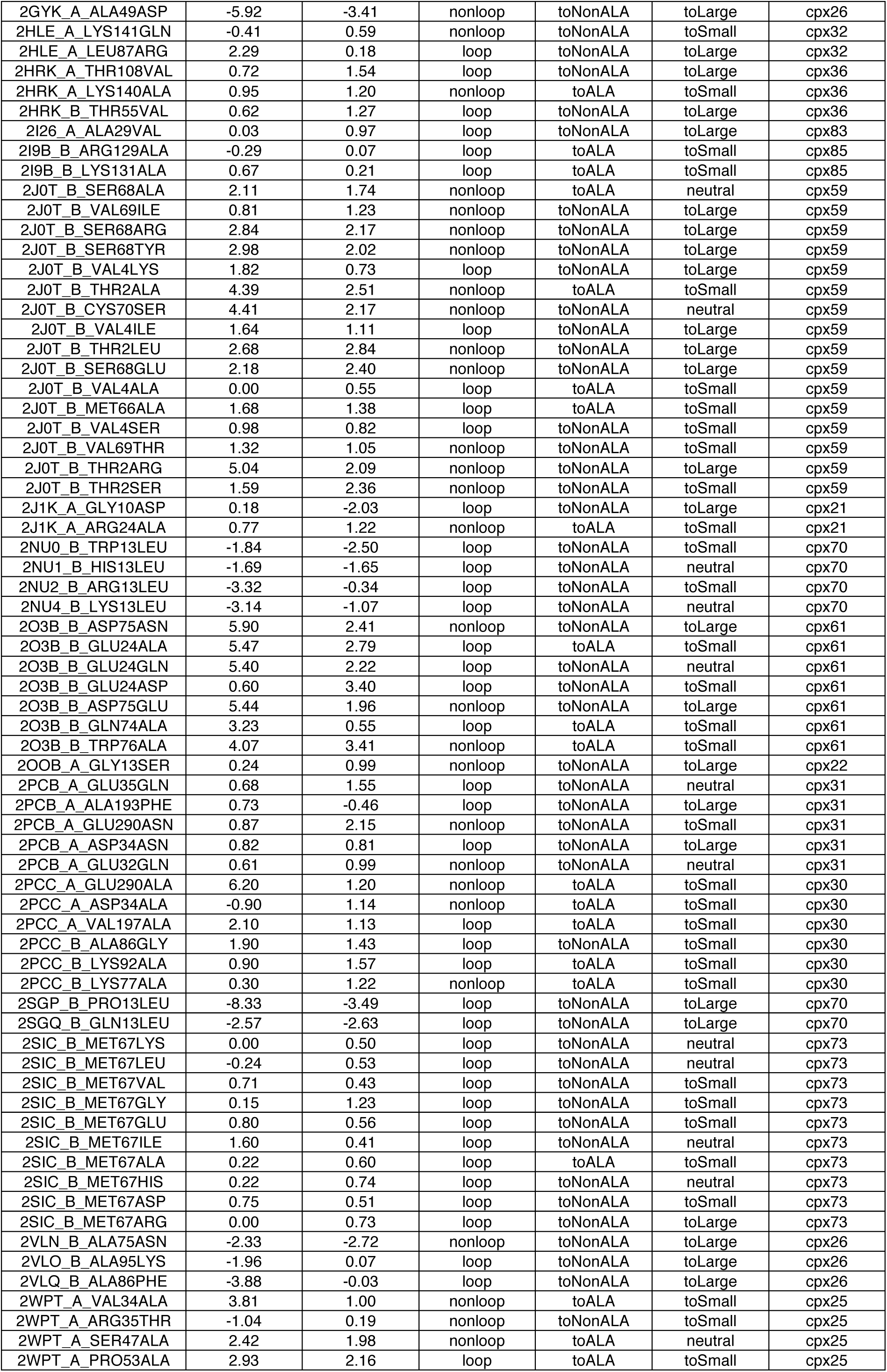
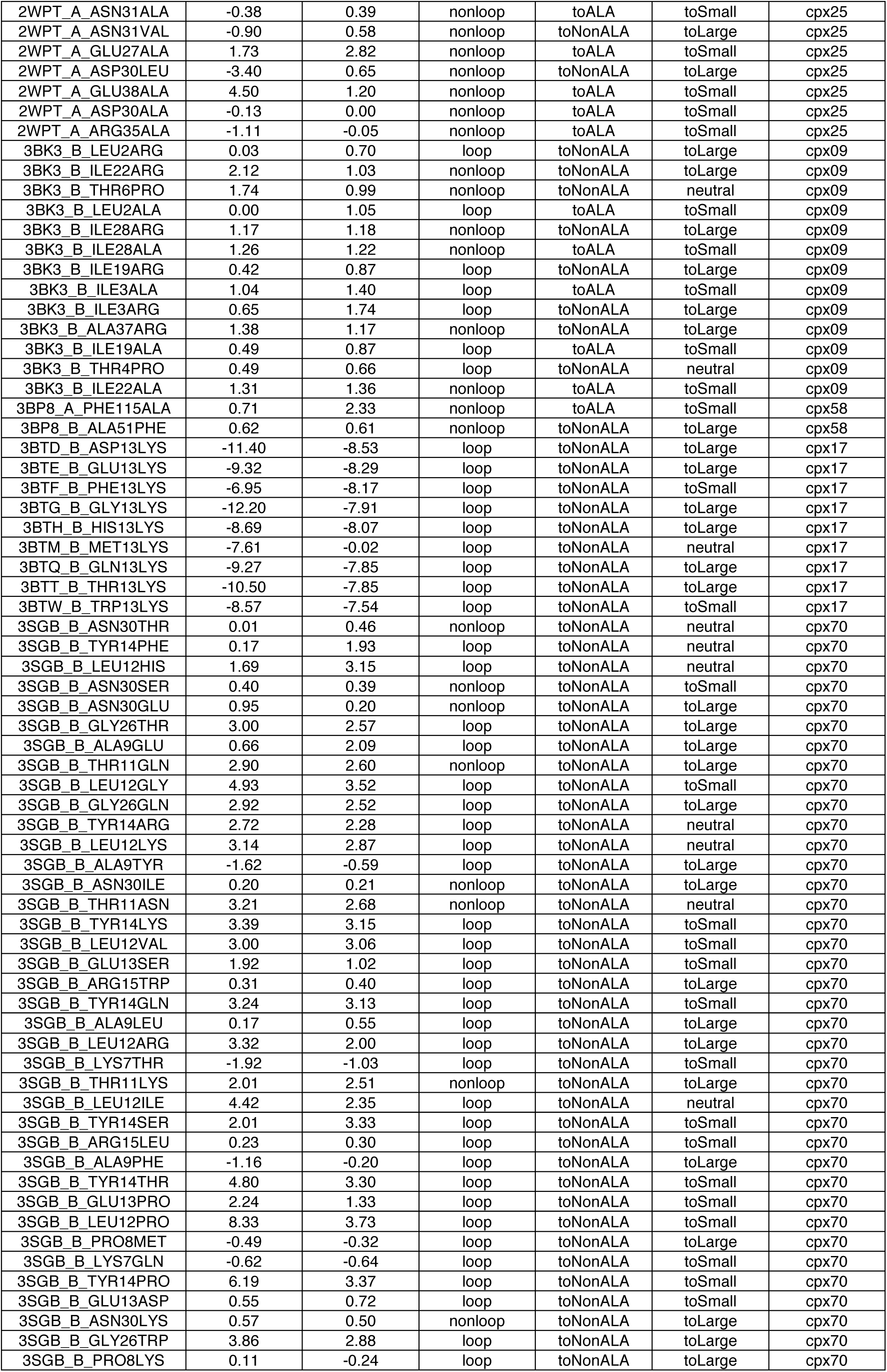
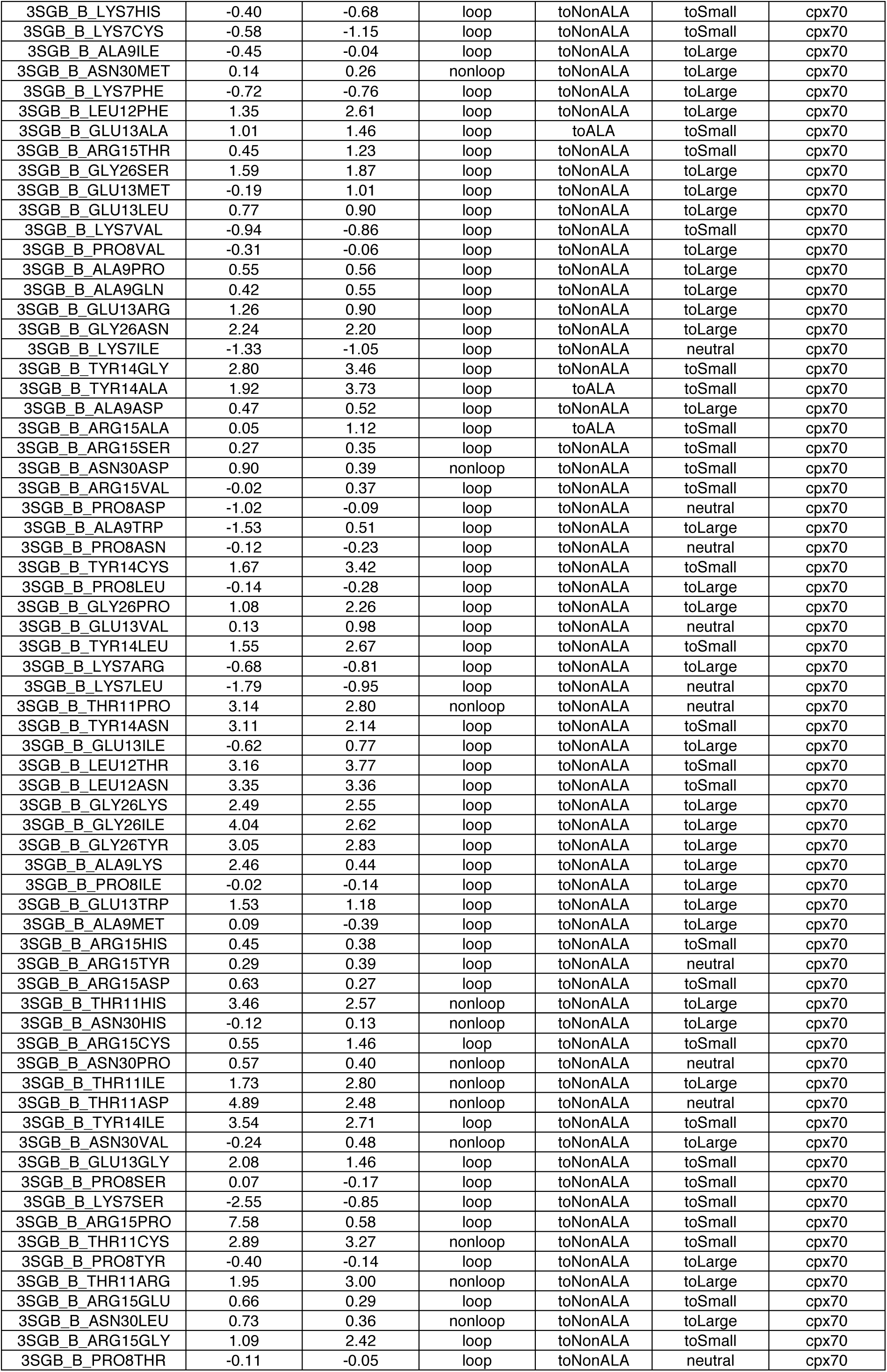
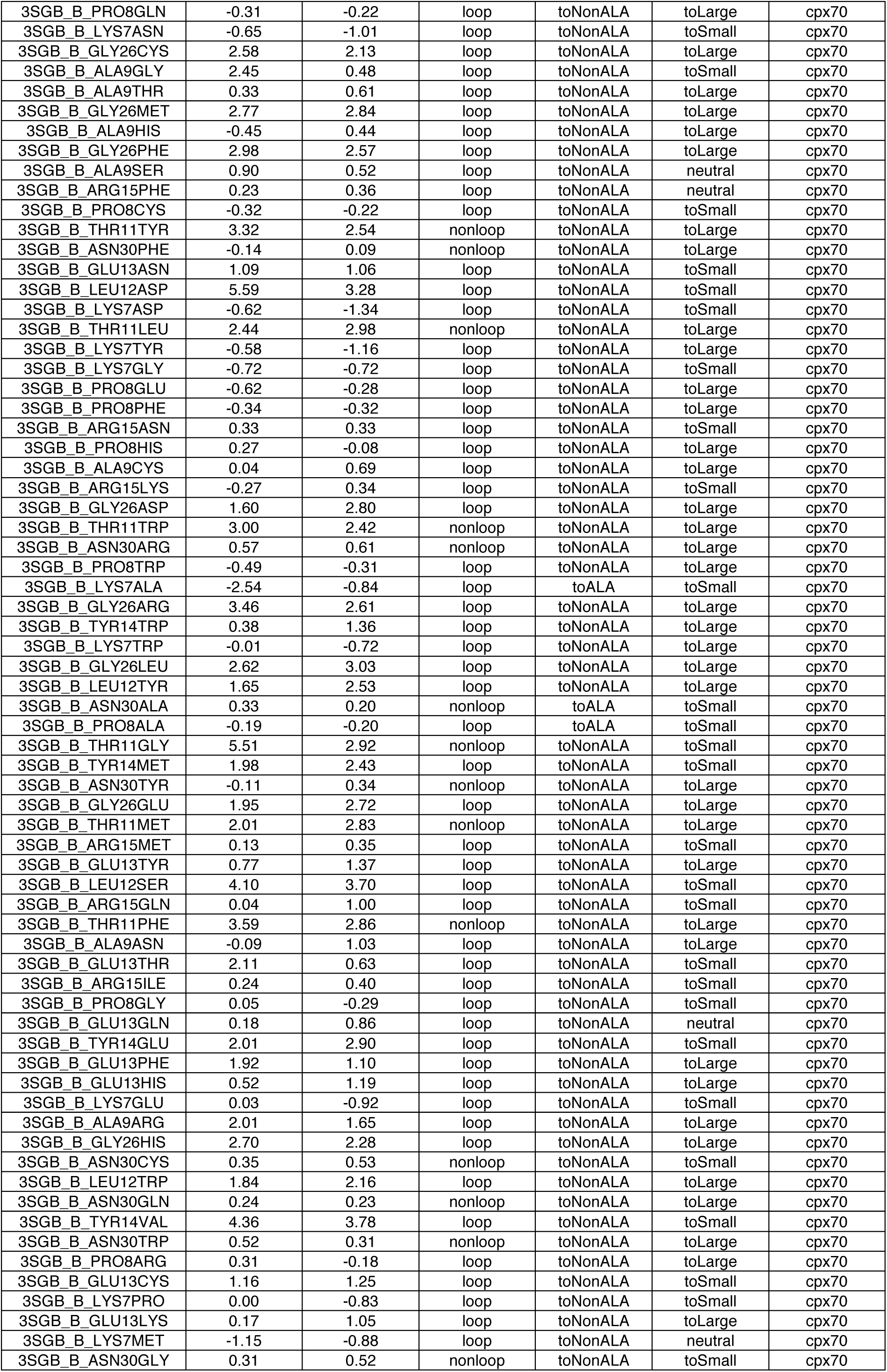
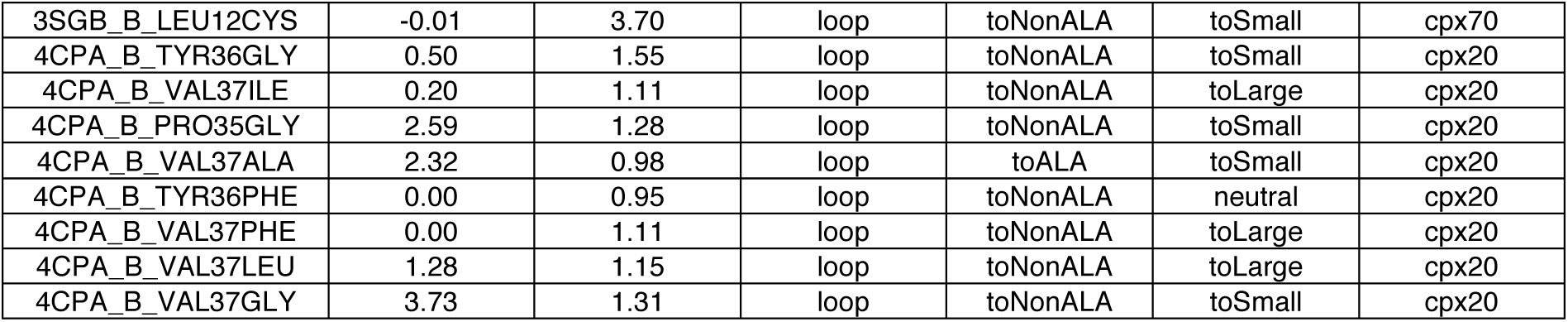
iSEE training dataset with experimental ΔΔG and average predictions of cross-validations. “SS” stands for the secondary structure (loop or nonloop), “muttype” for the type of mutated amino acid (toALA or toNonALA), “mutvol” for the change of amino acid size (toSmall, neutral, or toLarge), and “complex” for the index of complex types.

**Table S4.** NM test dataset with experimental ΔΔG and predictions of different methods. ΔΔG unit is kcal mol^-1^.

| mutationID | $\Delta\Delta G_{\text{exp}}$ | iSEE | pred1 | pred2 | CCPBSA | BeAtMuSiC | FoldX | mCSM | ZeMu | BindProfX |
| --- | --- | --- | --- | --- | --- | --- | --- | --- | --- | --- |
| 1IAR_B_ASN122ALA | 0.68 | 0.93 | 0.91 | 0.87 | 0.47 | 0.64 | 0.03 | 0.37 | -0.20 | 0.01 |
| 1IAR_B_ASP121ALA | 0.80 | 0.75 | 2.41 | 2.81 | 0.49 | 1.87 | 1.59 | 0.52 | 2.18 | 0.99 |
| 1IAR_B_ASP66ALA | 0.73 | 0.55 | 1.32 | 1.22 | 1.70 | 0.33 | 1.07 | 0.06 | 1.06 | 0.87 |
| 1IAR_B_ASP67ALA | 2.36 | 0.78 | 4.85 | 4.64 | 2.08 | 1.42 | 3.12 | 0.73 | 5.08 | 2.34 |
| 1IAR_B_ASP72ALA | 4.35 | 1.01 | 2.05 | 2.19 | 1.76 | 0.23 | 3.08 | 1.82 | 3.64 | 2.49 |
| 1IAR_B_LEU39ALA | 2.77 | 0.78 | 1.29 | 1.60 | 1.36 | 1.36 | 0.93 | -0.28 | 0.92 | 1.70 |
| 1IAR_B_LEU42ALA | 0.00 | 0.14 | 0.92 | 0.53 | 1.55 | -0.09 | 0.14 | 0.33 | 0.15 | 0.05 |
| 1IAR_B_LEU43ALA | 0.33 | 0.04 | 1.12 | 1.21 | 0.33 | 0.59 | 1.51 | 0.66 | 1.47 | 0.94 |
| 1IAR_B_PHE41ALA | 2.26 | 1.25 | 2.59 | 3.26 | 2.26 | 2.81 | 3.19 | 2.17 | 4.28 | 2.17 |
| 1IAR_B_SER70ALA | -0.24 | 0.16 | 0.96 | 1.22 | 0.62 | 1.41 | -0.46 | 1.14 | -1.26 | 0.82 |
| 1IAR_B_SER93ALA | -0.18 | 0.71 | 1.00 | 0.97 | 0.37 | 0.33 | 0.83 | 0.57 | -0.06 | 0.45 |
| 1IAR_B_TYR123ALA | 2.17 | 1.50 | 3.24 | 3.99 | 2.91 | 3.38 | 3.01 | 2.44 | 2.91 | 2.23 |
| 1IAR_B_TYR123PHE | -0.18 | 0.78 | 1.74 | 1.50 | 0.78 | 1.12 | -1.23 | 1.48 | -0.57 | 1.08 |
| 1IAR_B_TYR13ALA | 5.22 | 2.38 | 1.69 | 2.22 | 1.66 | 1.53 | 2.26 | 0.20 | 3.43 | 2.10 |
| 1IAR_B_TYR13PHE | 1.91 | 2.10 | 1.42 | 1.37 | 1.56 | 0.29 | 1.37 | 0.74 | 2.14 | 1.61 |
| 1IAR_B_TYR174ALA | 3.68 | 2.09 | 2.42 | 3.03 | 2.16 | 1.89 | 3.25 | 1.29 | 2.71 | 2.40 |
| 1IAR_B_TYR174PHE | 3.16 | 1.51 | 1.71 | 1.67 | 1.73 | 0.35 | 1.40 | 0.64 | 1.02 | 1.59 |
| 1IAR_B_VAL68ALA | 0.84 | 1.23 | 0.92 | 1.03 | 0.54 | 0.92 | 0.19 | 0.26 | -0.19 | 0.34 |
| 1IAR_B_VAL69ALA | 2.10 | 1.19 | 1.95 | 2.24 | 0.78 | 1.81 | 1.52 | 1.98 | 1.60 | 1.46 |

**Table S5.**
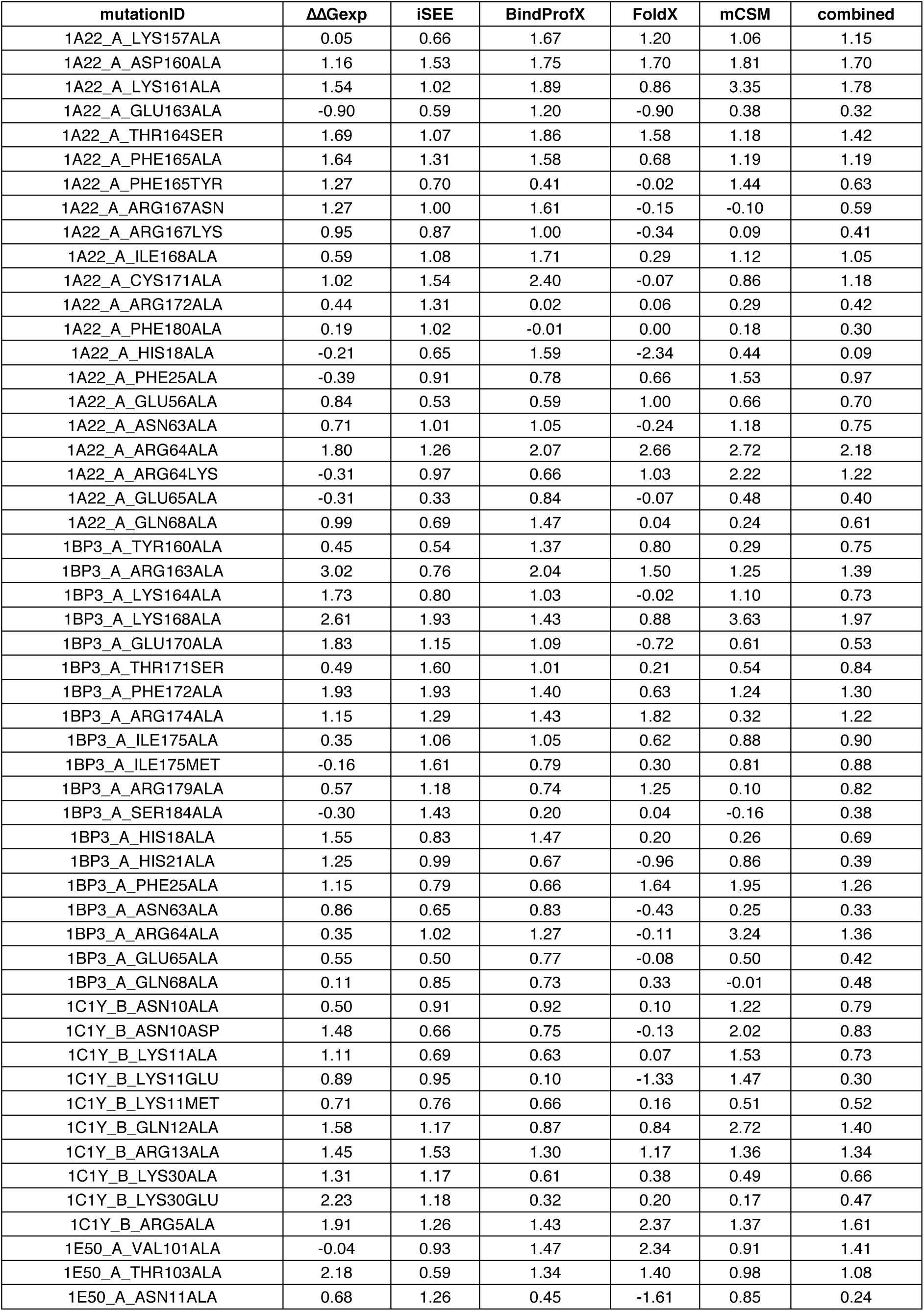
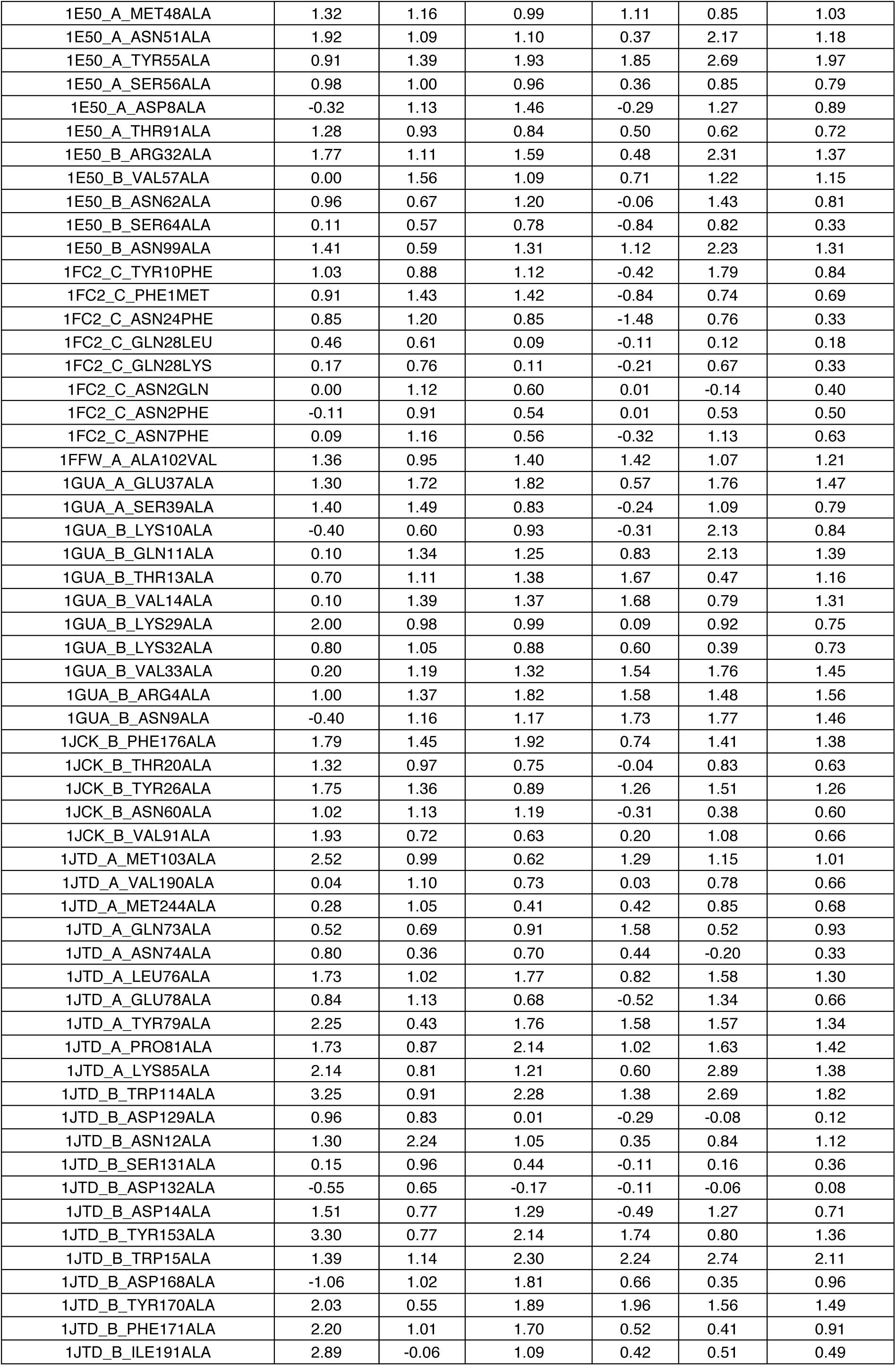
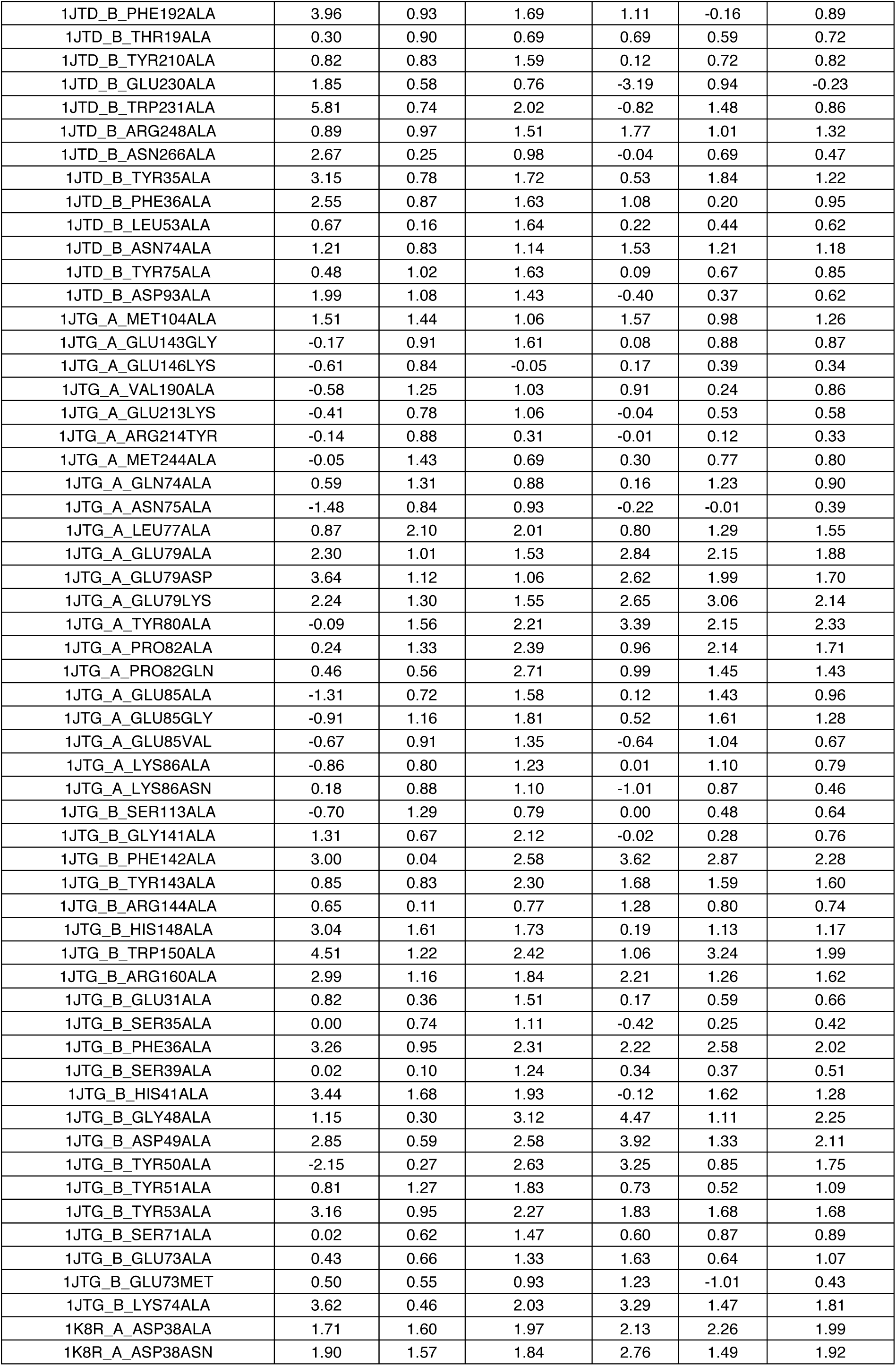
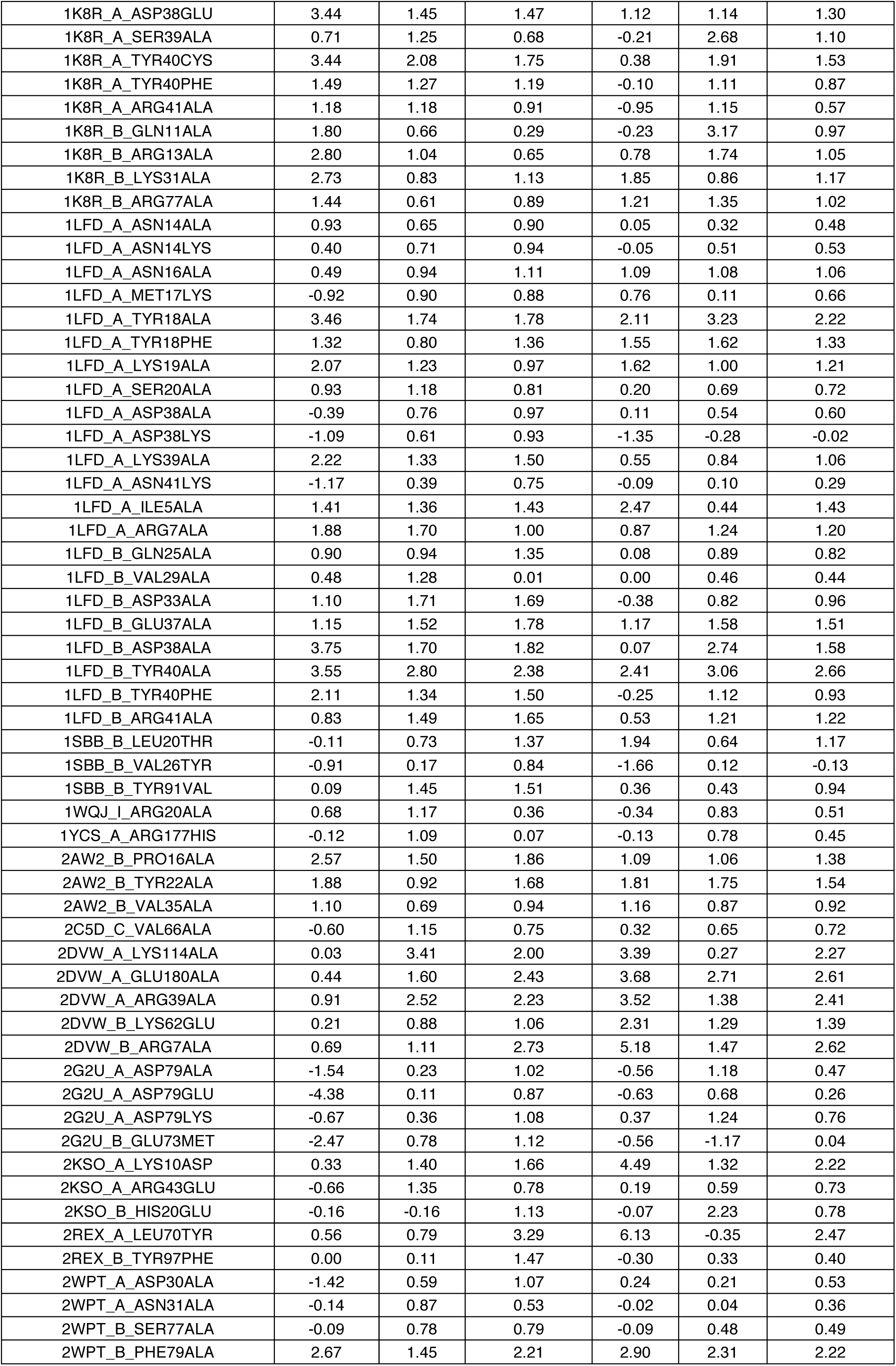
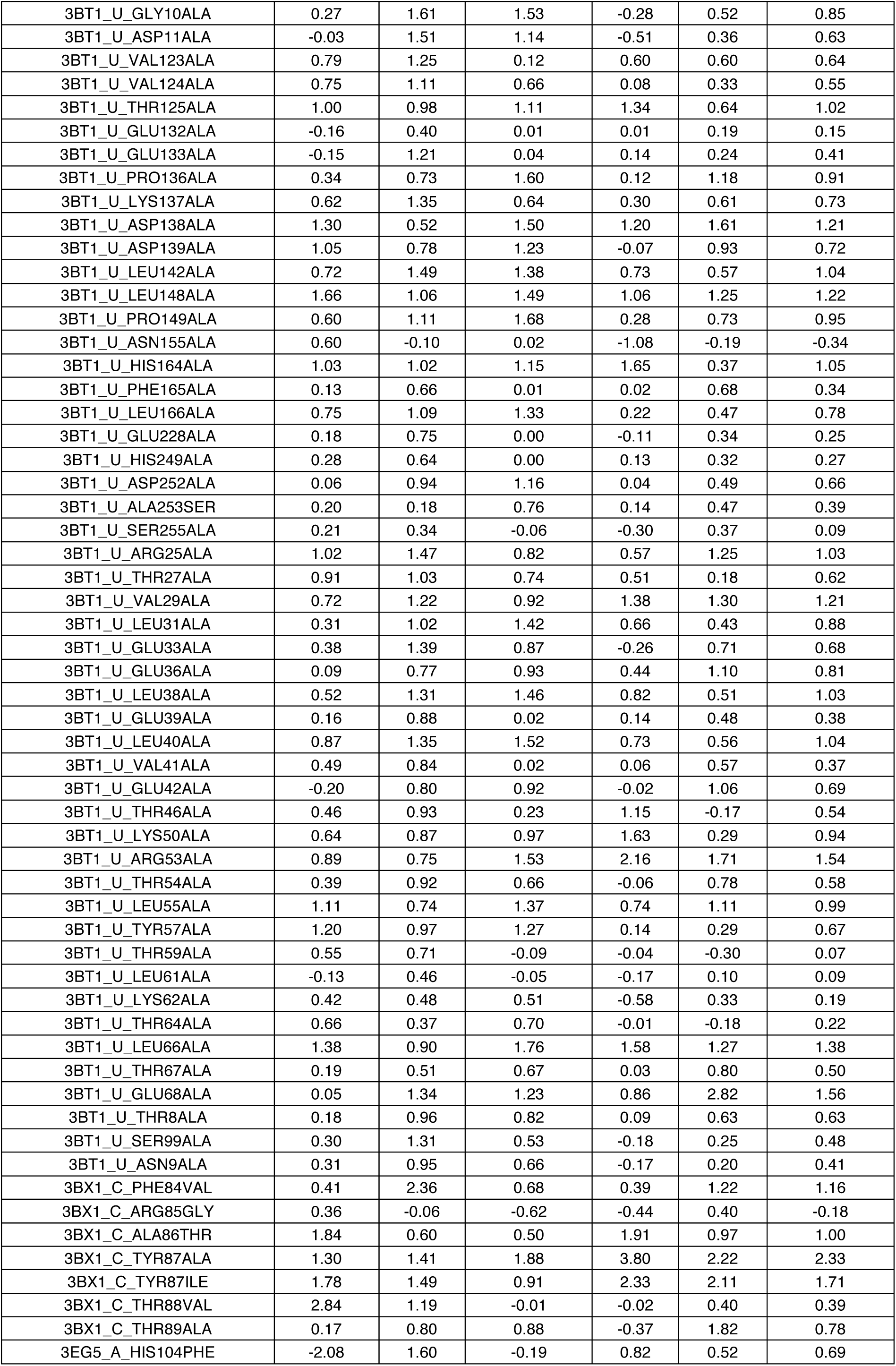
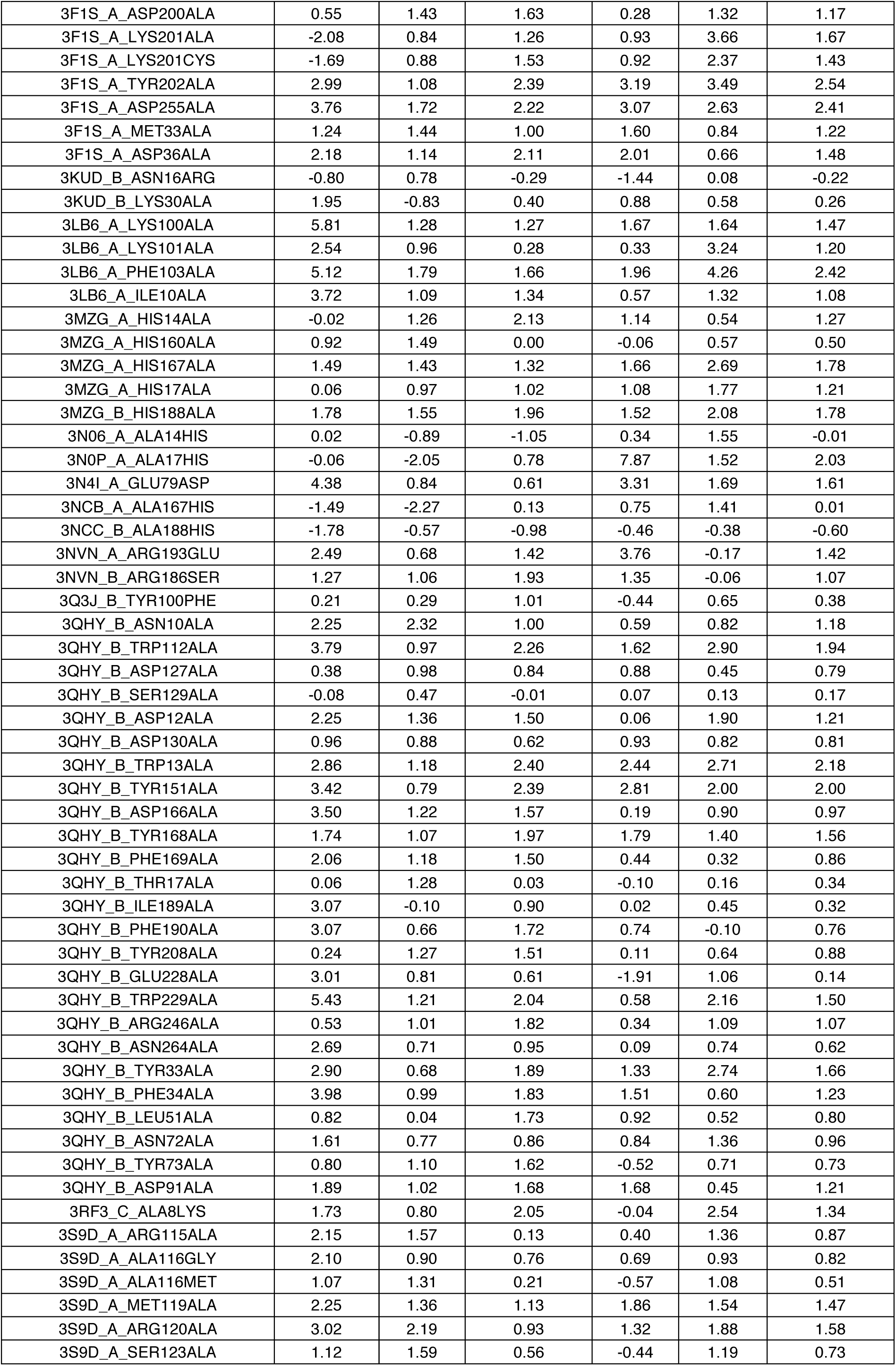
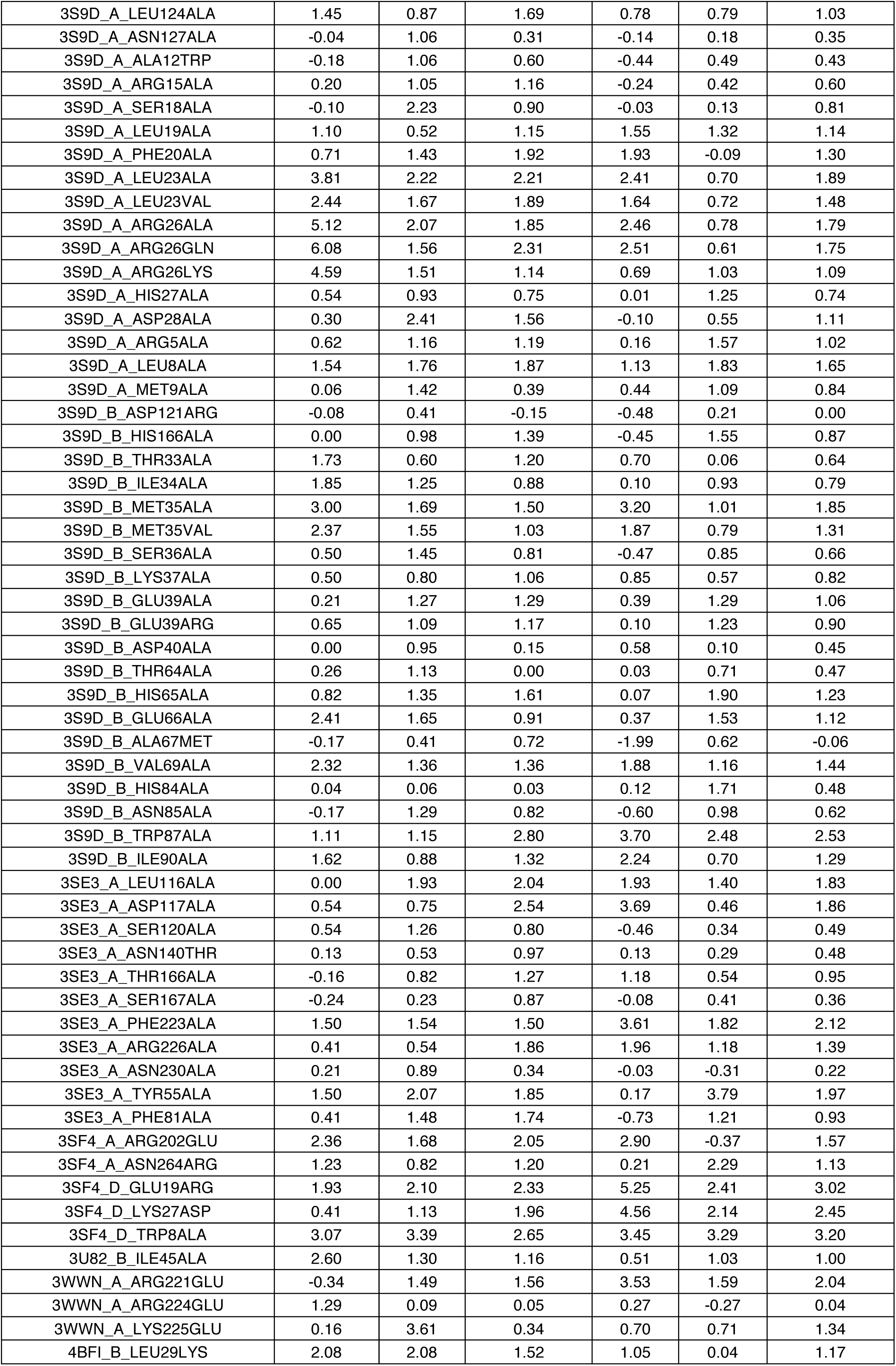
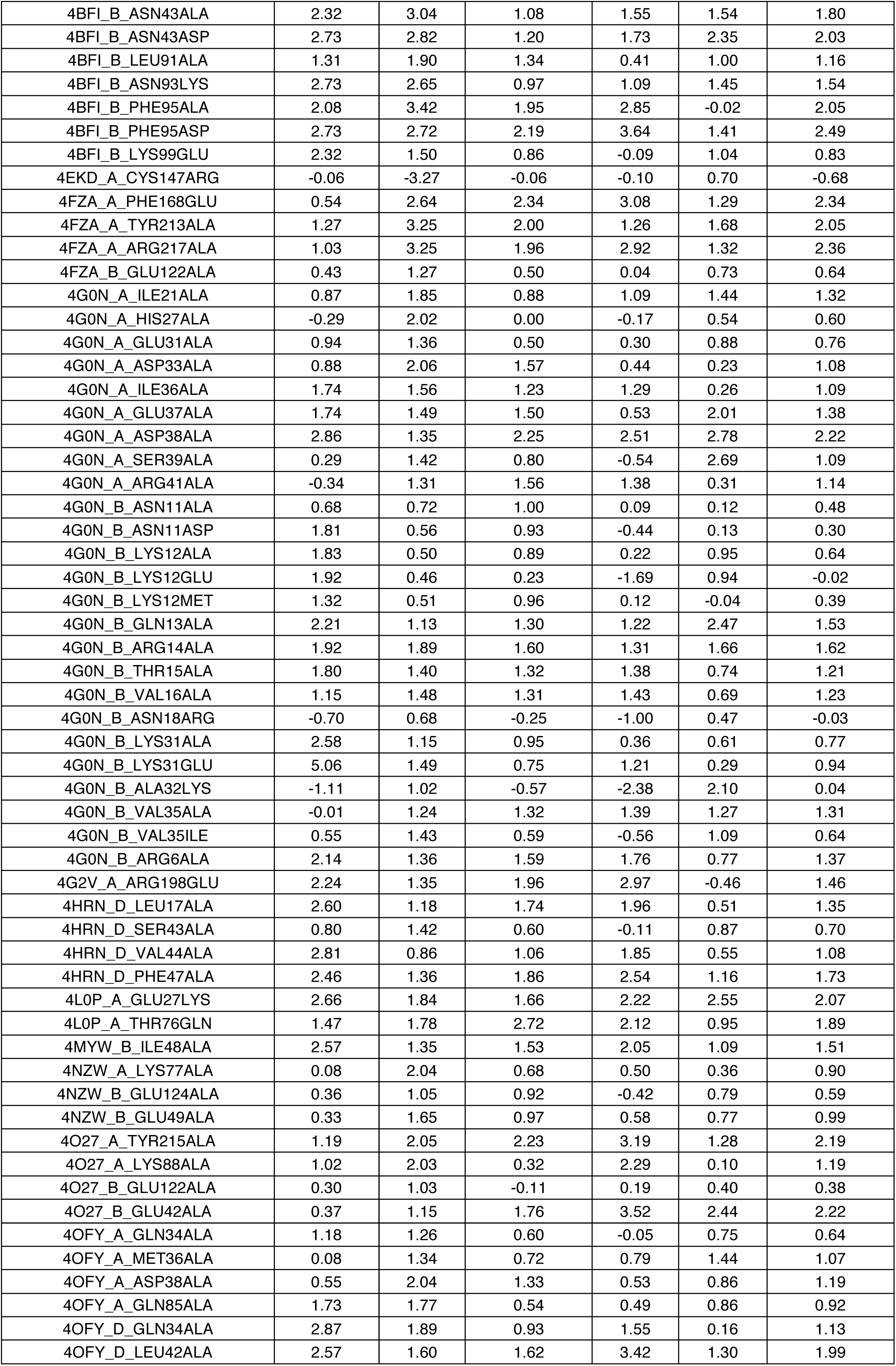
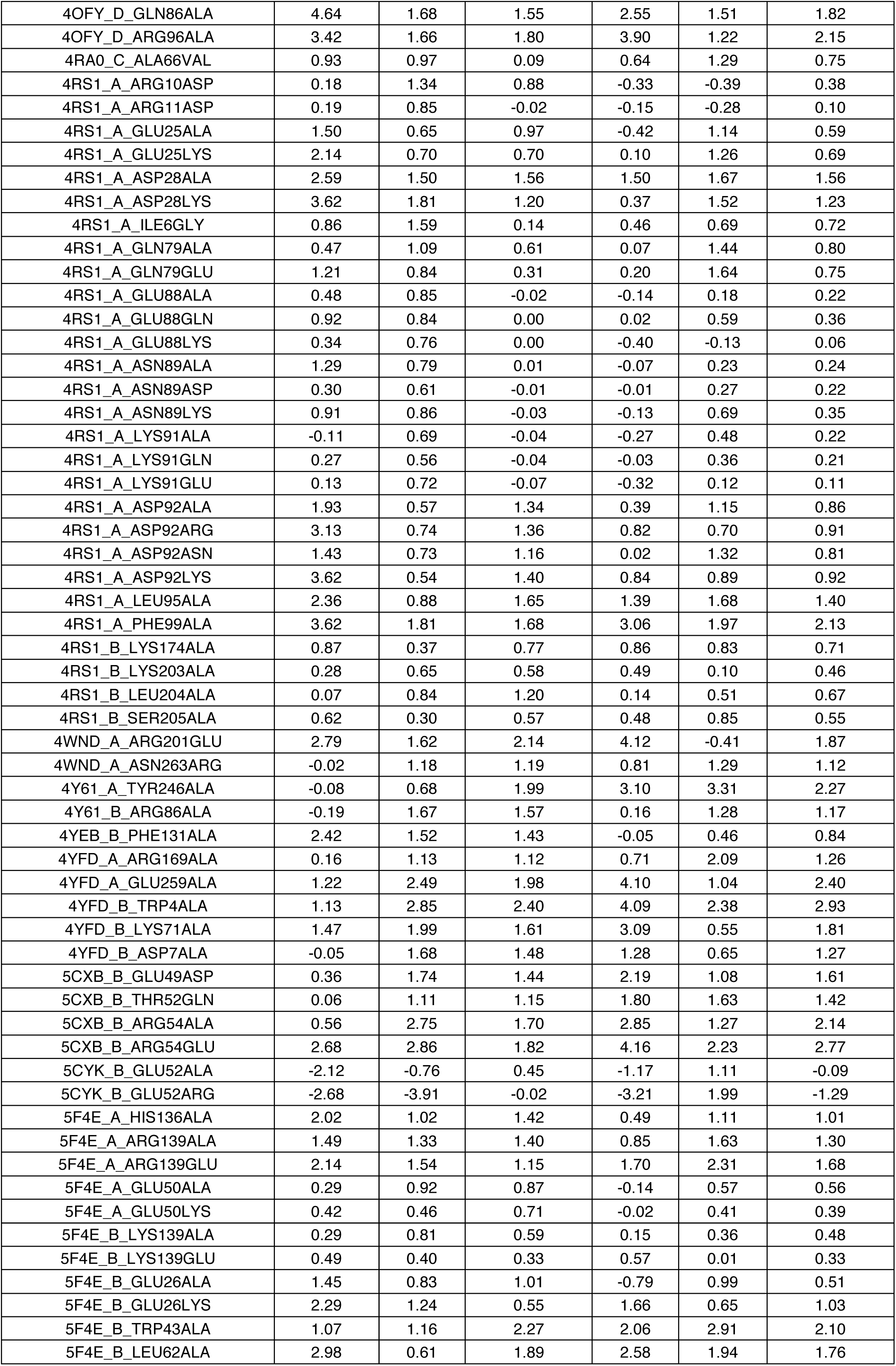
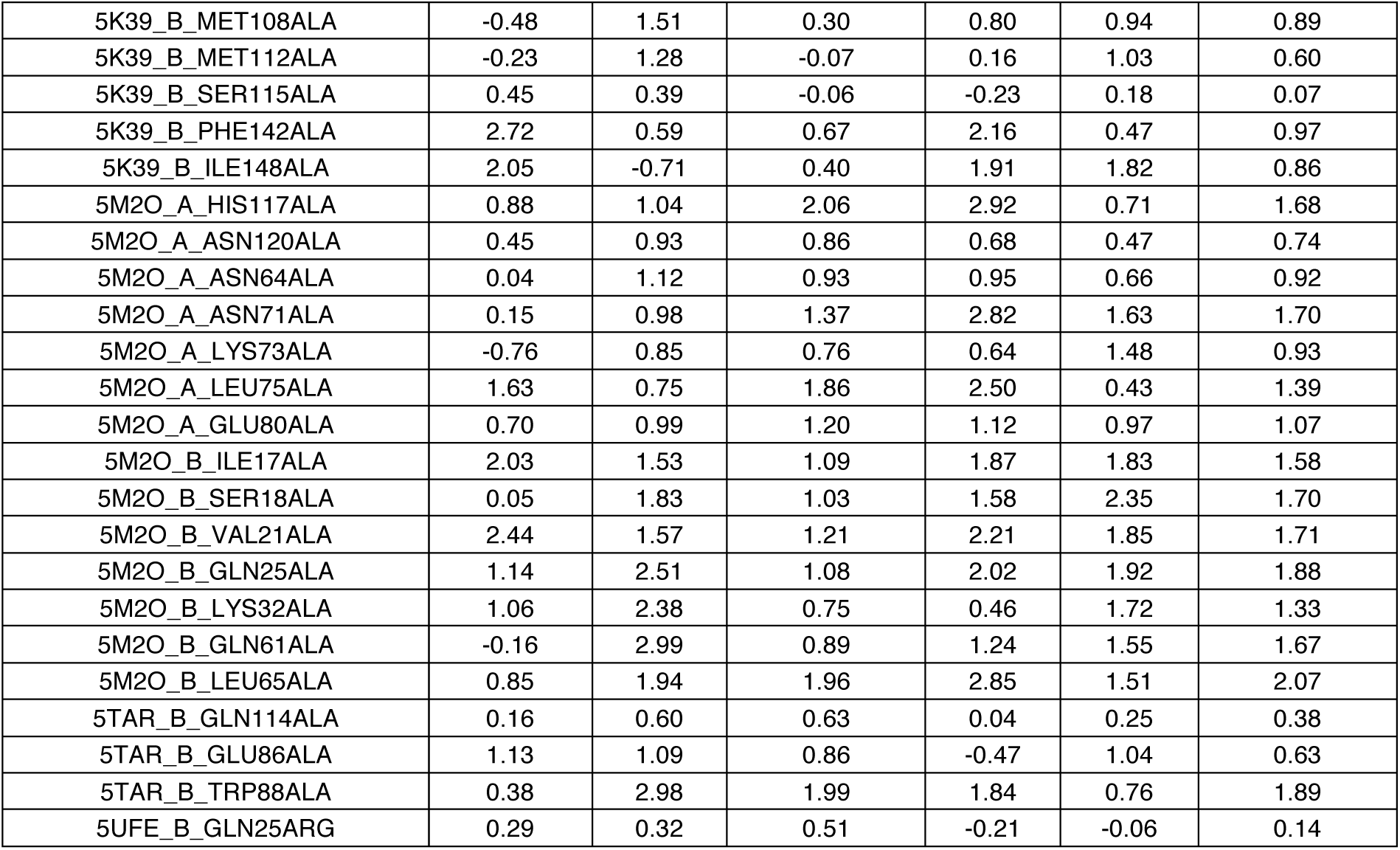
S540 test dataset with experimental ΔΔG and predictions of different methods. ΔΔG units are kcal mol^-1^.

